# Spatial and Temporal Resolution of Cyanobacterial Bloom Chemistry Reveals an Open-Ocean *Trichodesmium thiebautii* as a Talented Producer of Specialized Metabolites

**DOI:** 10.1101/2023.12.03.569781

**Authors:** Christopher W. Via, Laura Grauso, Kelly M. McManus, Riley D. Kirk, Andrew M. Kim, Eric A. Webb, Noelle A. Held, Mak A. Saito, Silvia Scarpato, Paul. V. Zimba, Peter. D. R. Moeller, Alfonso Mangoni, Matthew J. Bertin

## Abstract

While the ecological role that *Trichodesmium* sp. play in nitrogen fixation has been widely studied, little information is available on potential specialized metabolites that are associated with blooms and standing stock *Trichodesmium* colonies. While a collection of biological material from a *T. thiebautii* bloom event from North Padre Island, TX in 2014 indicated that this species was a prolific producer of chlorinated specialized metabolites, additional spatial and temporal resolution was needed. We have completed these metabolite comparison studies, detailed in the current report, utilizing LC-MS/MS-based molecular networking to visualize and annotate the specialized metabolite composition of these *Trichodesmium* blooms and colonies in the Gulf of Mexico (GoM) and other waters. Our results showed that *T. thiebautii* blooms and colonies found in the GoM have a remarkably consistent specialized metabolome. Additionally, we isolated and characterized one new macrocyclic compound from *T. thiebautii*, trichothilone A (**1**), which was also detected in three independent cultures of *T. erythraeu*m. Genome mining identified genes predicted to synthesize certain functional groups in the *T. thiebautii* metabolites. These results provoke intriguing questions of how these specialized metabolites affect *Trichodesmium* ecophysiology, symbioses with marine invertebrates, and niche development in the global oligotrophic ocean.

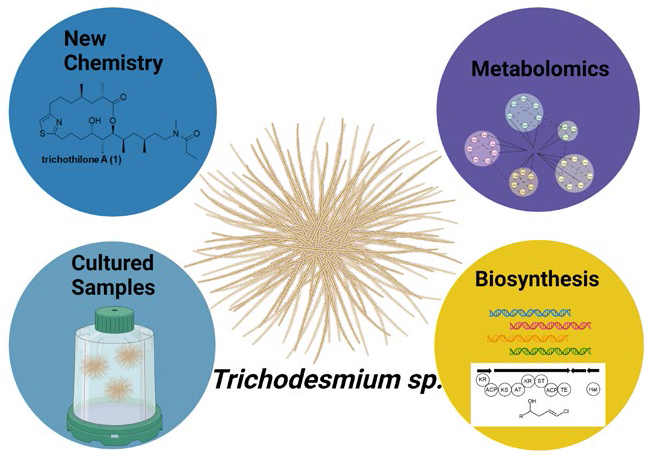

## Introduction

As climate change results in both increased global temperatures and CO_2_ concentrations in ocean waters, species of the cyanobacterial genus *Trichodesmium* are projected to expand their already substantial, oligotrophic ocean range as the subtropics move poleward.[1–3] An increase in oceanic and coastal growth of the periodically bloom-forming *Trichodesmium* will raise both the direct and indirect exposure risk to humans from potential toxins produced by this genus. While the ecological role of *Trichodesmium* as nitrogen fixers is well studied, the specialized metabolism of the genus along with the impact of potential toxins on human health and animal health are unknown. The latter is of increasing importance because of the aforementioned range and abundance expansions predicted for *Trichodesmium*.

Many aspects of *Trichodesmium*’s toxicity have been somewhat enigmatic and inconsistent. Multiple classes of toxins have been identified from *Trichodesmium* blooms and environmental collections. Previous *Trichodesmium* toxicity studies performed using homogenized cells, filtrates, aging cells, and crude extracts of *T. thiebautii* have shown toxicity to copepod grazers, while those from *Trichodesmium erythraeum* did not show the same levels of toxicity.[4,5] However, extracts of *T. erythraeum* samples obtained from coastal India showed toxicity to shrimp and multiple human cell lines.[6] It must be noted that species identifications in this study were done using morphology instead of genetic means, and no active component or components were structurally characterized.

Other research endeavors have isolated or detected individual metabolites, some with well-known toxicity from *Trichodesmium* spp. For example, saxitoxins and paralytic shellfish toxins have been identified from *Trichodesmium* blooms in the southwestern South Atlantic Ocean off the coast of Brazil,[7] in addition to samples collected in the coastal waters of India.[8] Hepatotoxic microcystins have been detected from *T. erythraeum* samples collected from waters off the coast of northwest Africa,[9] and from Indian waters in the Bay of Bengal.[8] The polyether toxin, palytoxin, was detected from *Trichodesmium* collected from a New Caledonian lagoon in the southwest Pacific Ocean.[10] The cyclic peptide trichamide was characterized from a cultured strain of *T. erythraeum* (IMS 101), but it was nontoxic.[11] A lipophilic ciguatoxin-like substance was isolated from *Trichodesmium* filaments and from Spanish mackerel specimens, *Scomberomorus commersoni*, collected from the same batches as the filaments, suggesting that *Trichodesmium* may be a vector for ciguatera poisoning and that fish may bio-accumulate toxins from blooms.[12] Focusing specifically on *Trichodesmium*, the many classes of toxins identified could result from either genomic differences in regionally specific strains, or from different environmental parameters modulating varied toxin profiles. However, an equally plausible scenario may be that biosynthesis is mediated by interactions between *Trichodesmium* and their associated organisms (collectively called *Trichodesmium* holobiont (TH) hereafter) or metabolite production by associated organisms. The TH is noteworthy for the diversity of organisms present (both in a bloom or colony) such as diatoms, copepods, dinoflagellates, heterotrophic bacteria, and other species of cyanobacteria [13–18]. Previous work has not included metabolite analysis of repeated field collections, biosynthetic gene cluster information, and culture confirmations, which has made interpreting toxicity studies challenging.

A key missing scientific investigation into potential *Trichodesmium* toxicity and specialized metabolism is an integrative, systematic study of the metabolites produced by *Trichodesmium* species in the environment and from isolates in culture. Our previously published work from a single large environmental collection showed that *T. thiebautii* dominated samples from the GoM produced potently cytotoxic compounds. We chemically characterized the components in extracts from a 2014 *Trichodesmium* bloom collected from the GoM. These efforts isolated and characterized several lipophilic, cytotoxic metabolites nearly all of which are hallmarked by a chlorovinylidene functionality. These metabolites represented new analogs from existing cytotoxic compound classes (e.g., smenamides C, D, and E),[19] and new classes of cytotoxic metabolites (e.g., the trichophycins, tricholides, and trichothiazoles), which had not been described previously.[20–23]

In the current report, we detail the results of continued field studies in 2017, 2019, and 2021 in the GoM to provide additional temporal and spatial resolution of specialized metabolite production. The culmination of these studies has shown that these metabolites can be detected in *T. thiebautii* colonies year after year and throughout the GoM. We have used genetic tools to determine the *Trichodesmium* species present in these collections and untargeted MS/MS-based molecular networking to provide a chemical inventory of colony and bloom metabolites. It appears clear from our data, that *T. thiebautii* in the oligotrophic GoM is a prolific producer of specialized metabolites, many of which possess nanomolar cytotoxicity against human cells (certain smenamides and smenothiazoles).[24,25] We report a new metabolite in this work, trichothilone A (**1**), which was confirmed in multiple *Trichodesmium* laboratory cultures. Additionally, we identified key genetic architecture in the *T. thiebautii* H94 genome that is consistent with the generation of key functional groups in many of the metabolites we have characterized, i.e., the chlorovinylidene group and terminal vinyl chloride present in the trichophycins and smenolactones, the trichotoxins, the smenamides, the conulothiazoles, and trichothiazole A.[19–21,23–30] This study provokes intriguing questions with respect to the full chemical potential of the members of this genus, and the potential for the development of metabologenomics approaches for the species in this group of ecologically important organisms.

## Results

### Morphological analysis and DNA sequencing of organism collections

Our previous work from an initial collection of *Trichodesmium thiebautii* biomass near North Padre Island, TX in 2014 provided the first evidence that there was a *Trichodesmium* species capable of prolific specialized metabolite production in the GoM. Mining this biomass resulted in the characterization of 24 new molecules,[20–23,26,29,30] including **1** presented in the current report. Additionally, seven chlorinated metabolites were detected from the 2014 collection via analysis of LC-MS/MS molecular networks. These metabolites were originally isolated and characterized from the sponge *Smenospongia aurea*, but putatively considered as the products of associated cyanobacteria due to the chlorovinylidene functional group present in these metabolites.[24,25,27–29]

The primary purpose of the current work was to determine if metabolite composition of these *T. thiebautii* colonies in the GoM was consistent over time and geographic location. To accomplish this, a second collection of *Trichodesmium* colonies was made in the GoM (27.00° N; 92.00° W) in 2017 (GoM2017, Table S1). Additionally in 2019, aboard the R/V Oregon II (National Oceanic and Atmospheric Administration), we visited several stations in the GoM and collected *Trichodesmium* colonies and biomass (GoM 2019-2-4, 6, 11, Table S1). Collections from 2017 and 2019 were identified as predominantly *Trichodesmium* sp. by examining filaments and puff and tuft colonies (Figure S1). Phylogenetic analysis of partial sequences of the 16S rRNA gene supported identification of samples as *T. thiebautii* (GoM2017, GoM2019-11), clustering with the *T. thiebautii* identified in the 2014 collection (Figure S2). During the 2019 research cruise, we collected bulk biomass from a surface accumulation (GoM2019-6) and were also able to harvest individual tuft and puff colonies from certain stations, sequestering individual colonies with a sterile loop and preserving them for laboratory analysis (e.g., GoM2019-11) (Figure S3). More field samples were collected from the GoM in 2021 (GoM2021-2-4, Table S1), and three cultivated samples of *T. erythraeum* IMS101 were examined - two separate cultures obtained from the Bigelow Laboratory for Ocean Sciences (IMS101) and another from the University of Southern California *Trichodesmium* Culture Collection (USCTCC) (*T. erythraeum* ST8) (Table S1). Finally, we examined legacy samples from a 2018 TriCoLim research expedition (AT39-05) in the Caribbean and Atlantic Ocean (TriCoLim St 3, 4, 6, 8, 13, 15, 16, 17, 20) (Table S1), which expanded the geographic coverage of metabolite composition of *Trichodesmium* colonies outside of the GoM. Collections and cultures used in this study are summarized in Table S1.

### LC-MS/MS-based molecular networking: spatial and temporal resolution of *Trichodesmium* metabolites

We utilized LC-MS/MS paired with molecular networking to compare the metabolite composition of *Trichodesmium* collections over time and geographic area in the GoM. The previously characterized metabolites from the 2014 collection served as a “screening library” for specialized metabolites (Figure S4). Collections from 2017 and 2019 showed remarkable consistency with respect to metabolite content when compared to the previous 2014 analysis. Several of the previously characterized metabolites from the 2014 collection were identified in samples from 2017 (21 metabolites annotated) and 2019 (18 metabolites annotated). These include many of the trichophycins, smenamides, tricholides, trichothiazoles, conulothiazoles, smenothiazole A, and trichothilone A (**1**) (Figure 1).

**Figure 1.**
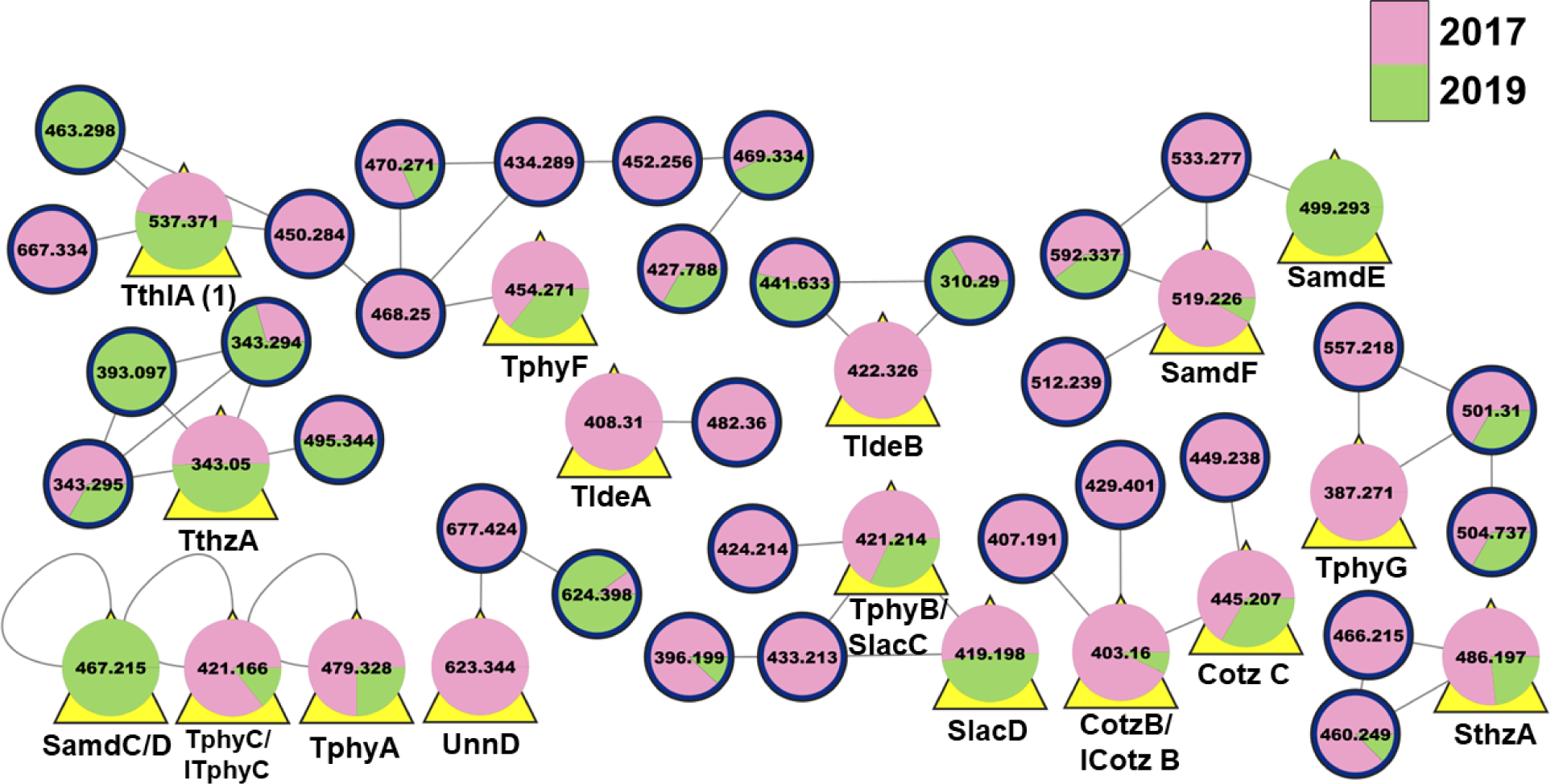
*T. thiebautii* specialized metabolites found year-after-year. Partial LC-MS/MS-based molecular network of extracts from collection GoM2017 (pink) and GoM2019-6 (green). Nodes are designated by their precursor masses and yellow triangles behind the node indicate that these metabolites were previously characterized by our group (cf. Figure S4). Cotz, conulothiazole; ICotz, isoconulothiazole; ITphy, isotrichophycin; Sam, smenamide; Slac, smenolactone; Sthz, smenothiazole; Tlde, tricholide; Tphy, trichophycin; Tthl, trichothilone; Tthz, trichothiazole; Unn, unnarmicin. The full network can be found at https://gnps.ucsd.edu/ProteoSAFe/status.jsp?task=2945b7552db5420dadd8e9167849d7e5.

Examining by geographic location using the 2019 samples, we again annotated many of the previously characterized metabolites from 2014 including trichophycins B and F, smenolactones C and D, smenamides A-F, isoconulothiazole B and conulothiazole C, trichothiazole A, tricholide B, and **1**. There were also over 25 metabolites that remain unidentified but were detected from all five stations (Figure 2).

**Figure 2.**
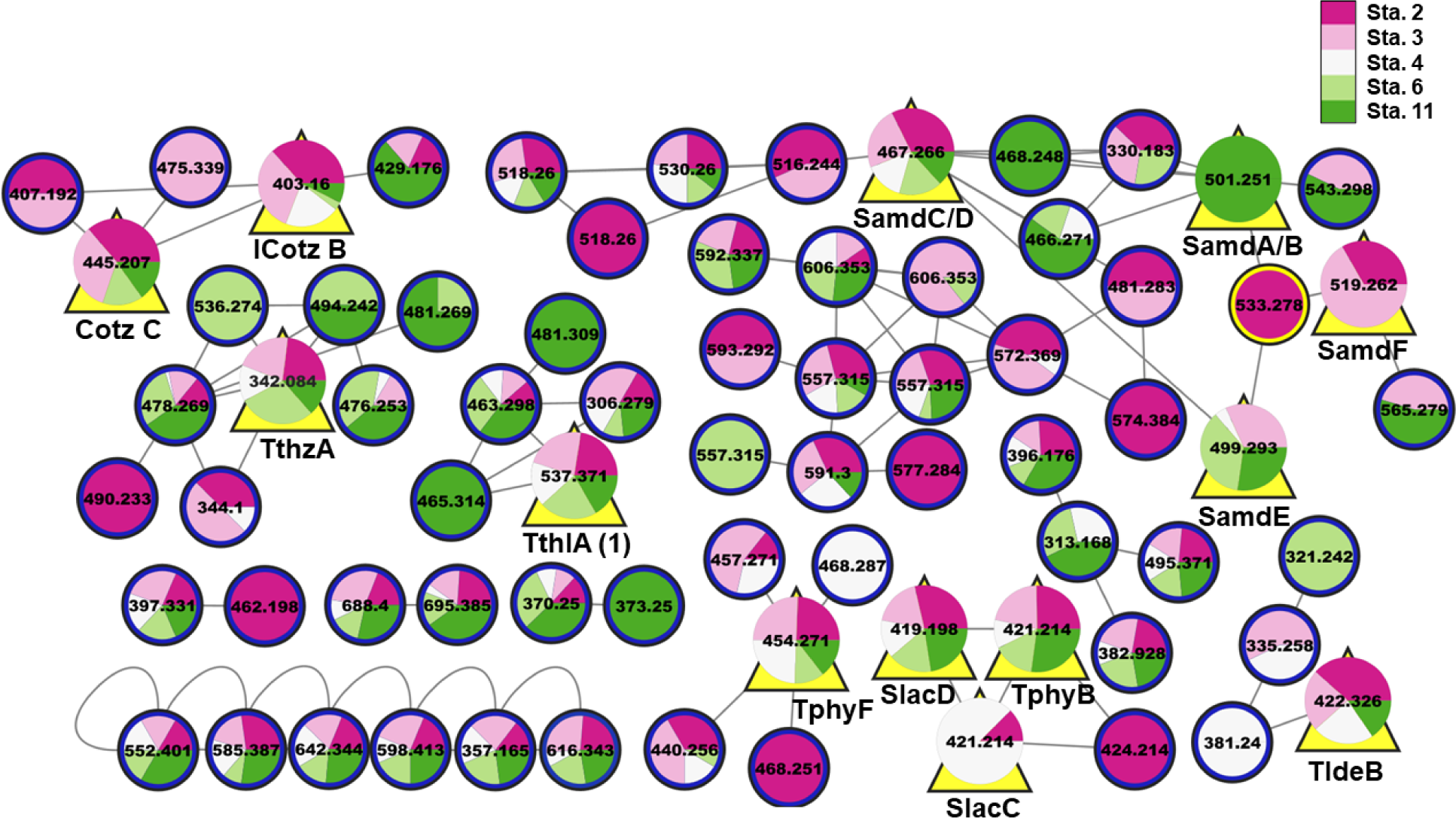
*T. thiebautii* specialized metabolites found across the GoM in 2019. Partial LC-MS/MS-based molecular network of extracts from 2019 collections (stations 2-4, 6, and 11). Nodes are designated by their precursor masses and “pie slices” designate stations where the metabolite was detected. Yellow triangles behind the node indicate that these metabolites were previously characterized by our group (cf. Figure S4). For abbreviations see Figure 1. The full network can be found at https://gnps.ucsd.edu/ProteoSAFe/status.jsp?task=52bda0f50dd94dd8b351f642e4d4598e.

We next analyzed the sample GoM2019-11, composed of ‘picked’ colonies (dominated by puff morphology). These colonies were collected via net tow, washed in a sieve, and transferred one-by-one to a preservation vial using a sterile loop. This sample does have associated bacteria in it, but it is devoid of other phytoplankton and loosely associated ectocommensals, which was verified by microscopy. Many of the previously characterized metabolites were identified in this sample as well (Figure 3). These results point to the existence of a core metabolome possessed by resident *T. thiebautii* colonies in the GoM, which has never been documented previously. This species produces hundreds of small organic metabolites, many of which are hallmarked by the incorporation of at least one halogen atom, typically chlorine (Figure S4). Following examination of all networks, one unidentified metabolite was prioritized for isolation and structure elucidation - *m/z* 537. This molecule was found in samples from 2014, 2017, all 2019 samples, and in the ‘picked colonies’ sample. Additionally, ion counts obtained from LC-MS/MS indicated that this molecule was abundant in all samples. In contrast to most *T. thiebautii* metabolites, mass spectrometry analysis indicated that **1** did not contain any halogen atoms. Following the metabolite composition analysis, HPLC-DAD- and LC-MS-guided isolation were performed.

**Figure 3.**
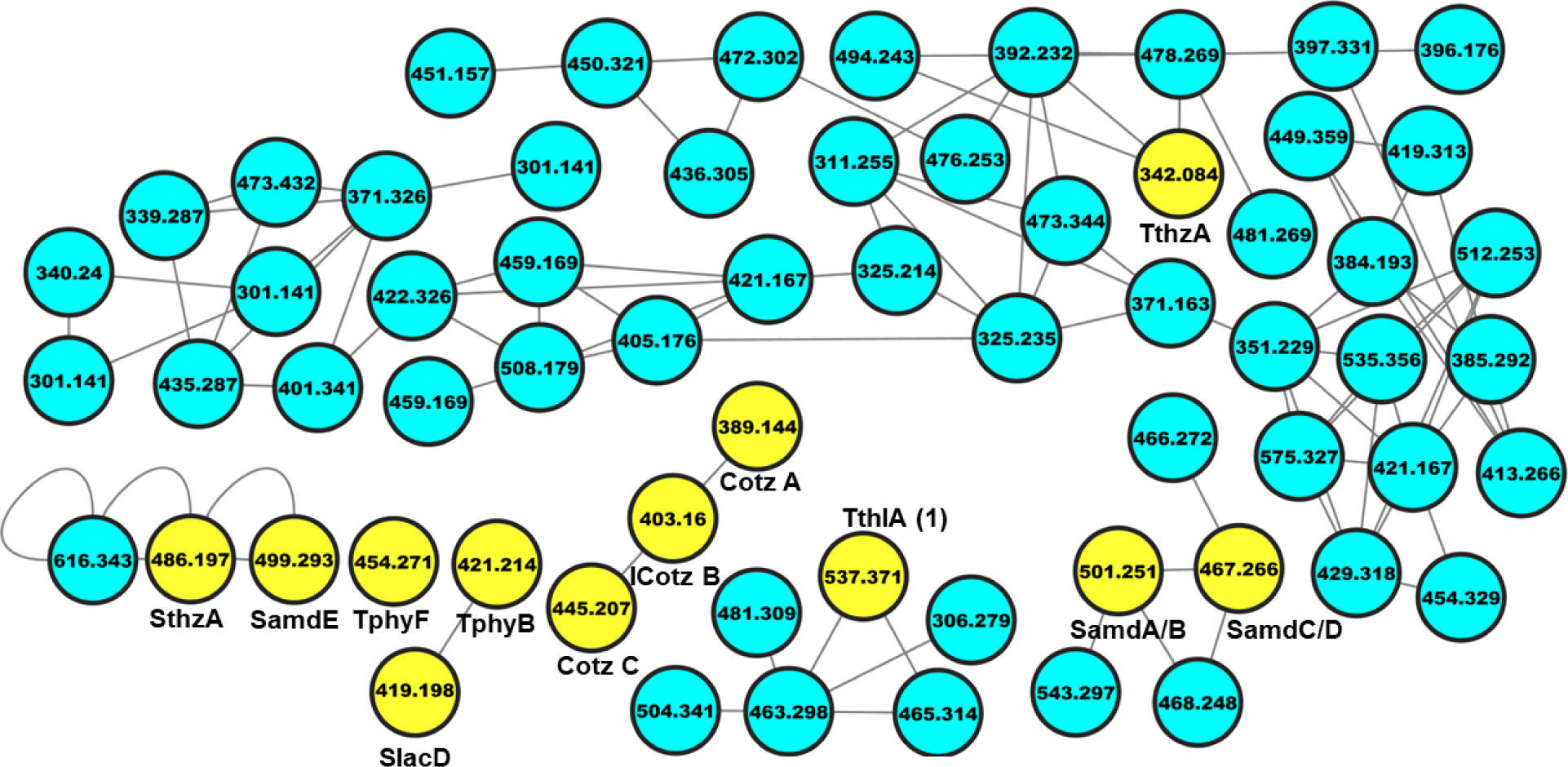
*T. thiebautii* specialized metabolites in “picked colonies”. Partial LC-MS/MS-based molecular network of extract from GoM2019-11. Colonies were hand-picked with a sterile loop from bulk collection and an extract was generated. Nodes are designated by their precursor masses. Yellow nodes indicate that these are metabolites were previously characterized by our group (cf. Figure S4). For abbreviations see Figure 1. The full network can be found at https://gnps.ucsd.edu/ProteoSAFe/status.jsp?task=838795d6f92a47db8ad23f3cb7c8bb6d.

### Structure characterization of 1

HPLC-DAD and mass spectrometry-guided isolation resulted in the isolation of an optically active pale yellow oil (**1**) (Figure 4A). HRESIMS analysis of **1** gave a protonated molecule [M+H]^+^ of *m/z* 537.3725 suggesting a molecular formula of C_30_H_52_N_2_O_4_S and a requirement of 6 degrees of unsaturation (Figure S5). Examination of 1D and 2D NMR data (Figures S6-S11) were used to establish the planar structure of the molecule. Certain ^1^H NMR resonances, most notably the *N*-methyl singlet, were split into two signals in a ratio of about 1:1. This suggested two conformers at the amide bond, which are detailed in Figure S12. The text discusses only the *Z*-conformer. This phenomenon has been described for other metabolites from *Trichodesmium* that contain an *N*-methyl amide functionality.[19] Inspection of the COSY spectrum and ^1^H-^1^H spin systems allowed for the assignment of three partial structures. The first from H-2 to H_2_-7 including the 1,3-dimethyl system from C-1 to C-5, and a second from H_2_-11 to H_2_-21 including a 1,3,5-trimethyl system from C-14 to C-20 (Figure 4B). A third partial structure consisted of a single COSY correlation between H_3_-30 and H_2_-29. HMBC correlations between H_2_-7 and C-8 (*δ*_C_ 156.0) and H_2_-11 and C-10 (*δ*_C_ 169.1) along with the H-9 resonance (*δ*_H_ 7.01) established a thiazole functionality that connected the first two partial structures. An *N*-methyl signal (*δ*_H_ 2.88) showed HMBC correlations to C-21 (*δ*_C_ 44.5) and C-28 (*δ*_C_ 172.1) and connected partial structure two to an *N*-methylpropanamide functionality. HMBC correlations of H-2 (*δ*_H_ 2.44) and H-16 (*δ*_H_ 4.81) with C-1 (*δ*_C_ 175.5) established the ester of a macrolactone and satisfied the final degree of unsaturation characterizing a hybrid polyketide-peptide metabolite that was named trichothilone A (**1**).

**Figure 4.**
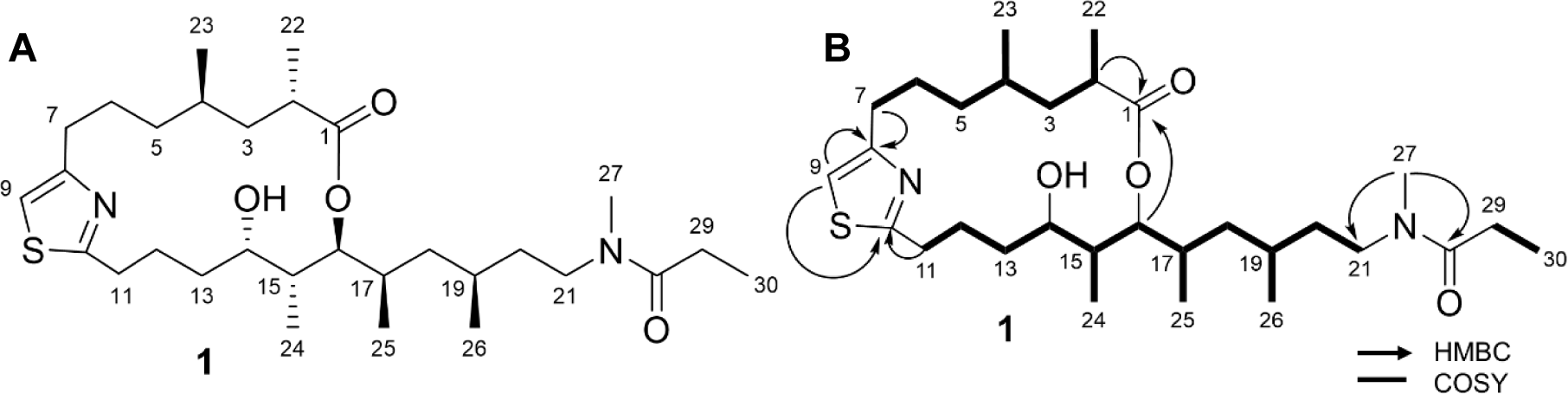
Structure of trichothilone A (**1**) (left) and key 2D NMR correlations (right).

**Table 1.**
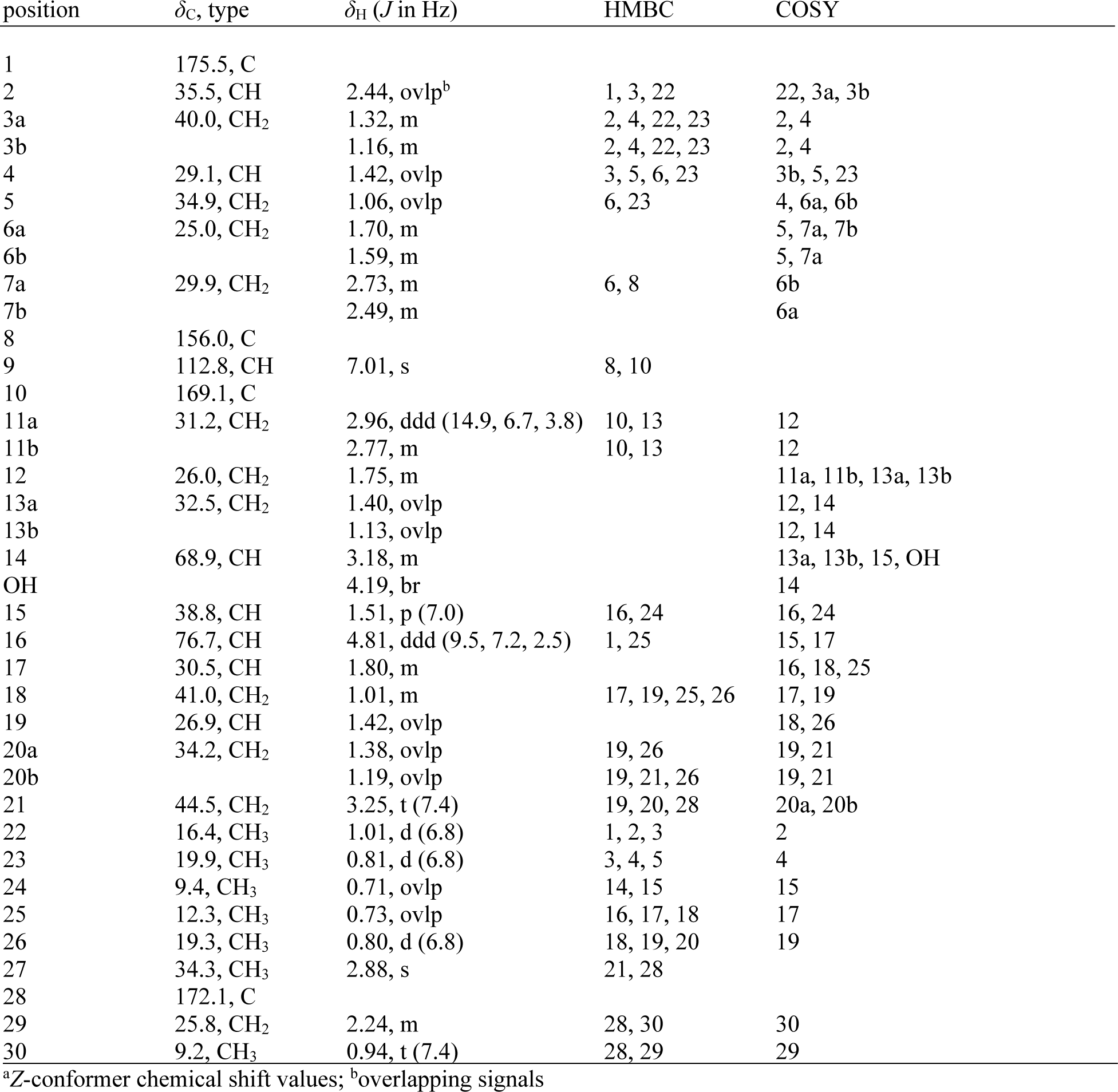
NMR data for trichothilone A (1)^a^ in DMSO-*d*_6_ (500 MHz for ^1^H NMR, 125 MHz for ^13^C NMR)

### Relative configuration of 1

Relative configuration of the four stereogenic centers, C-14 to C-17 was determined as 14*S*\*,15*R*\*,16*S*\*,17*R*\* using the Murata’s approach,[31] which is based on ^1^H-^1^H and ^1^H-^13^C scalar couplings and NOE effects, as detailed in Figure S13. The ^1^H-^13^C scalar couplings were directly measured using a HECADE-HSQC experiment (Figure S13),[32] or estimated as “large” or “small” from the ratio of the relative magnitudes of their cross peak measured in the HMBC spectrum with respect to a common proton, an intense peak indicating a large ^2,3^*J*_CH_, a weak or missing peak a small ^2,3^*J*_CH_.[33] Further support to the configuration of the segment C-14 to C-17 of **1** was provided by Kishi’s method for the relative configuration of contiguous propionate units.[34,35] Comparison of ^13^C NMR signals in **1** recorded in three different solvents (CDCl_3_, CD_3_OD, and DMSO-*d*_6_) to Kishi’s ^13^C NMR database determined the relative configuration as α,α,β,β for this segment (cf. Table S2 and Figure S14), fully confirming the stereochemical assignment based on Murata’s method.

Determination of the relative configuration at C-2 and C-4 in **1** with respect to the C-14/C-17 segment was achieved using quantum-mechanical computational chemistry, namely DFT prediction of ^1^H and ^13^C NMR chemical shifts (DFT-NMR).[36] Considering the structural complexity of compound **1**, the truncated model compound **1m** was used for the calculations, in which the flexible side chain of the natural compound is replaced by an isopropyl group. This strongly reduced the number of low-energy conformations of the molecule, while it did not significantly affect the chemical shifts of the region of the molecule under study.[37] The four possible stereoisomers of **1m** at C-2 and C-4 (*RR-***1m**, *RS-***1m**, *SR-***1m**, *SS-***1m**) were considered in the calculations (Figure 5).

**Figure 5.**
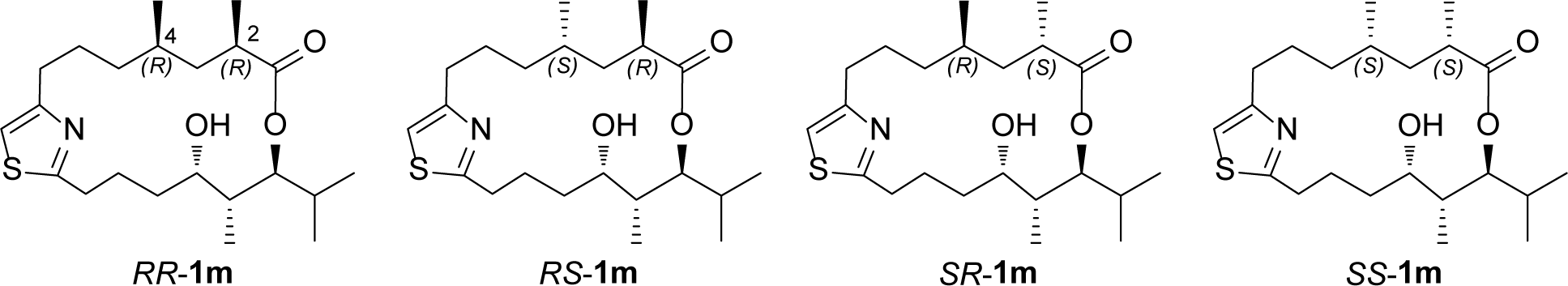
The four possible stereoisomers of the model compound **1m** at C-2 and C-4 (*RR-***1m**, *RS-***1m**, *SR-***1m**, *SS-***1m**) considered in the DFT calculations.

Conformational analysis of the four **4m** stereoisomers was performed using the program Pcmodel 10.0 (GMMX algorithm, MMFF94 force field),[38] which generated 35 through 66 conformers for each stereoisomer within 4 kcal/mol from the lowest-energy conformer (see SI Experimental Procedures for details). All the conformers were optimized by DFT minimization using the Gaussian program at the B3LYP/6-31G(d,p)/SMD level,[39] and their ^1^H and ^13^C chemical shifts were calculated at the mPW1PW91/6-311+G(d,p)/PCM level. Average chemical shifts over all the conformers were then calculated for each of *RR-***1m**, *RS-***1m**, *SR-***1m**, *SS-***1m** based on the Boltzmann distribution, and compared with the experimental values measured for **1**. Comparison of the root-mean-square deviations (RMSD) of ^13^C and ^1^H chemical shifts immediately showed that *SR-***1m** fit experimental data much better than the other three stereoisomers (*RR-***1m**: 2.32 and 0.150 ppm; *RS-***1m**: 2.24 and 0.145 ppm; *SR-***1m**: 1.98 and 0.122 ppm; *SS-***1m**: 2.46 and 0.149 ppm, respectively). The DP4+ statistical analysis of computational results fully confirmed this,[40] in that it assigned 100.00% probability to *SR-***1m** being the correct stereoisomer (Figure S15). The *anti* orientation of the methyl groups at CH_3_-22 and CH_3_-23 of **1** determined above was further supported by the Δ(H_a_−H_b_) value (0.16 ppm) of the methylene protons attached to C-3, which is consistent with the trend shown for 1,3-*anti* dimethyl groups in the macrocycle myxovirescin (*anti* = 0.20 ppm; *syn* = 0.70 ppm).[41] Therefore, the relative configuration of **1** was extended to 2*S*\*,4*R*\*,14*S*\*,15*R*\*,16*S*\*,17*R*\*, and only configuration at C-19 remained to be determined.

An attempt to determine the relative configuration at C-19 using DFT-NMR was unsuccessful because the predicted chemical shifts of the two possible epimers showed similar agreement with experimental data, with DP4+ probability <80% and therefore not conclusive (data not shown). Moreover, the Δ(H_a_−H_b_) value of the methylene protons attached to C-18, which was close to 0, could not be used for determination of relative configuration of the 1,3-dimethyl system CH_3_-25/CH_3_-26 because the method is unreliable when the methyl branches are adjacent to a lactone, which is the case for this system in **1**.[41] To overcome this limitation, we generated the acyclic methyl ester of **1**, characterized this derivative (**2**) using HRMS and ^1^HNMR (Figures S16 and S17), and reanalyzed this compound to determine the chemical shifts of the intervening methylene unit in the 1,3-dimethyl system. The Δ(H_a_-H_b_) value of the methylene protons attached to C-18 of **2** (Δ(H_a_-H_b_) = 0.43) strongly supported a *syn* relative configuration for CH_3_-25 and CH_3_-26 (Figure S14).[41] In addition, the Δ(H_a_-H_b_) measured for **2** was almost identical to the value (0.40 ppm) measured for the methylene protons attached to C-5 of the 4-bromophenacyl derivative of hemibourgeanic acid (**3**),[42] thus ruling out the possibility that the stereogenic centers C-16 and C-15 could affect the reliability of the method (Figure 6). These data defined the full relative configuration of **1** as 2*S*\*,4*R*\*,14*S*\*,15*R*\*,16*S*\*,17*R*\*,19*S*\*.

**Figure 6.**
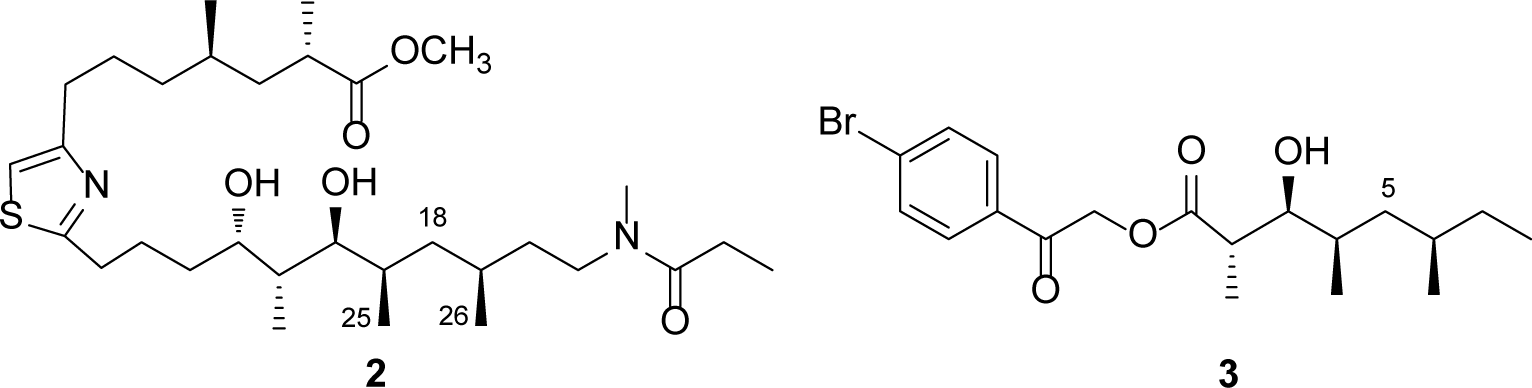
Comparison of Δ(H_a_-H_b_) values for the methylene protons attached C-18 in **2** (0.43) and the Δ(H_a_-H_b_) values for those protons attached to C-5 (0.40) in the model compound hemibourgeanic acid (**3**).[42]

### Absolute configuration analysis of 1

A single secondary alcohol in **1** (attached to C-14) allowed us to generate Mosher esters at that position. We analyzed the *S* and *R* esters of **1** via ^1^H NMR, COSY, and TOCSY. A positive Δ(*δ*_H_*S*-*δ*_H_*R*) value for H-14 and negative Δ(*δ*_H_*S*-*δ*_H_*R*) values for H-9 and H-11 supported a 14*S* configuration (Figure S18). Relaying stereochemical assignments based on the previously determined relative configuration supported a 2*S*,4*R*,14*S*,15*R*,16*S*,17*R*,19*S* configuration of **1**.

### Cytotoxicity of 1

Trichothilone A (**1**) showed modest cytotoxicity against neuro-2A cells with an IC_50_ value of 13.7 µM (Figure S19). Due to the limited amount remaining following chemical degradation and derivative formation experiments, we were not able to investigate other bioactivities that the molecule might possess.

### Specialized metabolites in cultures and from collections in the wider Caribbean and Atlantic Ocean

Trichothilone A (**1**) was detected in both the cultivated *T. erythraeum* IMS101 samples and the *T. erythraeum* ST8 sample with matches to retention time, high resolution mass spectrometry measurement and MS/MS fragmentation pattern (Figure S20). An examination of the cell mass and media of a second IMS101 culture also showed the presence of trichothilone A (**1**) (Figure S21), and its presence in the media may be related to its ecological relevance. Examining collections from the GoM collected aboard NOAA’s R/V Oregon II in 2021 (revisiting stations from 2019) and samples from the TriCoLim expedition,[43] we continued to identify the presence of metabolites from our 2014 library by matching MS/MS fragmentation patterns. The GoM samples showed the presence of certain trichophycins, smenamides, and other molecules (Table S3). We detected trichothiazole A in extracts from TriCoLim stations 3, 8, 16, and 17 (Figure S22) and trichophycin F in 3, 6, and 17 (Figure S23). We also detected smenamide A/B from station 3 extracts (Figure S24).

### Genomic analysis of *T. thiebautii* H94, *T. erythraeum* IMS101, and field metagenomes

We examined several genome and metagenome assembled genomes (MAGs) that were publicly available,[43] searching for genetic architecture consistent with the unique functional groups (chlorovinylidene group and terminal vinyl chloride) present in *T. thiebautii* metabolites (although these functional groups were not in **1**). We also examined genomes and MAGs from *T. thiebautii* and *T. erythraeum* for putative full biosynthetic gene clusters (BGCs) that might include **1**. Specifically, this included the genomes of *T. thiebautii* H94, *T. erythraeum* IMS101, and the metagenome assemblies obtained from samples collected during the R/V Atlantis TriCoLim cruise,[43] having some colocation with the archived samples from the cruise that we analyzed chemically during this work. Analysis of PKS clusters present in the *T. thiebautii* H94 genome using the antiSMASH platform[44] and the known cluster BLAST function identified a module in the *T. thiebautii* H94 genome which showed similarity to modules in the curacin A pathway.[45] Curacin A possesses a terminal alkene functionality and the biochemistry surrounding its construction and the activity of the decarboxylating thioesterase has been experimentally shown.[46] Additional mining and annotation using DELTA-BLAST[47] showed the presence of the following domains in the *T. thiebautii* cluster of interest: ketosynthase (KS), acyltransferase (AT), ketoreductase (KR), acyl carrier protein (ACP), sulfotransferase (ST), and a thioesterase domain (TE). These domains are consistent with those that form the terminal alkene in curacin A.[45] Furthermore, adjacent to this module was a gene that BLAST searching annotated as an Fe(II)-dependent halogenase. This partial cluster was similar to the genes predicted to form the terminal vinyl chloride in many of the trichophycins and trichothiazole (Figure S25A). A second partial gene cluster showed remarkable consistency with the genetic architecture necessary to form the chlorovinylidene group. An HCS cassette was present in this cluster consisting of an HMG-CoA synthase and two enoyl CoA hydratases and ketosynthase, acyltransferase, and three ACP domains with additional enoyl reductase and ketosynthase domains (Figure S25B). The HCS cassette has been observed in gene clusters that produce products with the chlorovinylidene moiety, such as the jamaicamide biosynthetic pathway.[48] While a specific halogenase was not identified in this partial cluster, the elements are present for the vinylidene group observed in nearly all the metabolites isolated from *T. thiebautii* collections (Figure S4). The HMG-CoA synthase was also identified in MAGs Bin1_Station19 and Bin1_Station18, which phylogenomics analysis supported as *T. thiebautii*.[43]

Investigations of the assemblies of *T. erythraeum* IMS101 and the MAG assemblies from the field samples did not show these partial clusters described above but a different PKS-NRPS gene cluster was identified in *T. erythraeum* IMS101 and several of the field samples (Bin3_Station18_1, *T. erythraeum*; Bin4_Station16, *T. erythraeum*; Bin5_Station11, *T. erythraeum*; and Bin8_Station4_1, *T. erythraeum*), which is of interest (Figure S26). In addition to the core PKS-NRPS architecture in this cluster, a lipocalin-like gene was annotated, which has been seen in the malyngamide pathway and other pathways and is hypothesized to be involved in cyclization.[49] However, we could not unequivocally connect this pathway to a specific product including **1** and further biochemical investigations will be necessary to characterize this pathway. We did identify elements of the trichamide gene cluster[11] in *T. thiebautii* H94, Bin4_Station 16 (*T. erythraeum*), Bin3_17_1 (ND), Bin2_Station 16 (*T. thiebautii*) Bin2_Station14_1 (ND), Bin2_Station5_1 (ND), Bin1_Station19 (*T. thiebautii*) Bin1_Station18 (*T. thiebautii*), Bin1_Station16_1 (ND), and Bin1_Station15_1 (*T. thiebautii*) showing its widespread presence in *Trichodesmium*.

## Discussion

### *Trichodesmium* species as toxin and specialized metabolite producers

With the occurrence of cyanobacterial blooms increasing concurrent with climatological changes, there is need to determine the ecological role of specialized metabolites and how they affect speciation, ecophysiology, and range expansion. The rarity of *Trichodesmium* cultures stems from an inability for most laboratories to obtain hand samples and to scale-up cultures of *Trichodesmium* for the biomass amounts that can lead to the isolation of metabolites. This has necessitated that many toxin and specialized metabolite isolation efforts come from environmental collections. While many classes of toxins have been detected from these blooms, it is difficult to attribute production of these metabolites, including **1**, to *Trichodesmium* without analysis of axenic cultures or complete biosynthetic gene cluster information. However, the widespread presence of **1** in environmental collections and its detection in three independent unialgal cultures of *Trichodesmium* supports its cyanobacterial production, and its presence in the media may indicate an important ecological role. The *T. thiebautii* dominated blooms and collected colonies showed intriguing profiles with respect to their metabolite composition. While our laboratory has characterized several new specialized metabolites from a *T. thiebautii* dominated bloom collected near North Padre Island, TX, this is the first report of the confirmation of these molecules from subsequent collections across space and time. Continuing longitudinal and geographic studies will aid in understanding the metabolite profiles of this group and potentially lead to understanding the ecological role of these metabolites. The widespread presence of trichothiazole, trichothilone A (**1**), and other metabolites presumes they likely have an important physiological or ecological role. We have established a demonstrated analytical capability to show that certain specialized metabolites are found in colonies in the GoM and the wider Caribbean and Atlantic Ocean, but *Trichodesmium* is found around the globe with notable repeated blooms in the Pacific and Indian Oceans.[50] It would be most interesting to continue analysis in collections from these areas using our established methods to provide further foundational studies on *Trichodesmium* specialized metabolism. The ability to accumulate more material from yearly blooms will enhance the potential for **1** and other metabolites to be comprehensively evaluated for their potential therapeutic benefit and ecological function. Furthermore, there remain hundreds of uncharacterized metabolites as shown in the various molecular networks we have generated. We are only scratching the surface of the molecular potential of *Trichodesmium*.

### Connecting biosynthetic pathways to *Trichodesmium* specialized metabolites

A current limitation in our results is the lack of identification of complete *T. thiebautii* putative biosynthetic gene clusters for the metabolites we have characterized, and a complete understanding of the species assemblages in environmental samples. Field samples are more accurately defined in a clade format dominated by one of the two major groups of *Trichodesmium* (*T. erythraeum* or *T. thiebautii*) and recent work has shown that there are four clades of N_2_-fixing *Trichodesmium* (*T. erythraeum* A & B, *T. thiebautii* A & B).[43] Until there is an unequivocal biosynthetic identification of complete gene clusters in the *T. thiebautii* genomes, and the lack of these clusters in strains of *T. erythraeum*, no definitive answer can be given on species chemotaxonomy and metabologenomics. There is also previously published evidence that indicates environment populations are comprised of mixed species.[51] Defining species assemblages in collections will be necessary to understand community composition and how this may affect specialized metabolite production. Furthermore, while the cultivated sample of *T. erythraeum* only showed the presence of one of the specialized metabolites we have characterized (**1**), the culture conditions may not be conducive to metabolite production, or we do not understand the environmental triggers necessary for metabolite expression. Examination of the publicly available genome of *T. erythraeum* IMS101 did not unequivocally identify any biosynthetic gene clusters as the likely architecture to produce **1** or other specialized metabolites. However, many intriguing elements were present in the *T. thiebautii* H94 genome and other MAGs as mentioned above with respect to the terminal vinyl chloride and chlorovinylidene groups, and there are additional partial clusters that are consistent with the biosynthesis of characterized metabolites. For instance, there are multiple NRPS modules in Bin1_Station 18 that are consistent with the biosynthesis of the smenothiazoles and conulothiazoles including an adenylation domain for cysteine and a heterocyclization domain that likely would generate the thiazole functionality and NRPS module predicted to incorporate valine and proline, which are consistent with the constitution of smenothiazole A. While dozens of molecules have been characterized from *Trichodesmium* by our group, they have similar carbon scaffolds, and most are analogs of each other. We speculate that the biosynthesis of these products in *Trichodesmium* follows that deciphered for the vatiamides in which a unique combinatorial non-collinear hybrid PKS-NRPS system generated a series of analogs adding to molecular diversity.[52]

Holistic molecular approaches must be taken to define species assemblages, genomic analysis to define biosynthetic architecture, and culture and field experiments to understand potential abiotic and biotic triggers that control metabolite production. Additionally, as the bulk of these samples were from field collections, there are many variables such as the impact of the bacterial and microbial community and the sample volumes collected vary widely from a few colonies to bulk collections. Instituting water volume measures, colony counts, and more molecular community data will aid in better detection and more comparable data across samples. Furthermore, efforts will include creating sensitive and selective targeted mass spectrometry methods to assay for these metabolites in samples from around the world. Triple quadrupole mass spectrometers using multiple reaction monitoring, screening known compound-specific fragments, can detect hundreds of metabolites and the development and implementation of this approach will create an efficient workflow for assessing field samples and cultures in a more targeted manner. Additionally, integrating other omics approaches with metabolomics, such proteomics[53] has the potential to reveal the processes responsible for *Trichodesmium* ecological success. Recent work has shown that along the West Florida Shelf there is a coastal vs. open ocean separation for *T. erythraeum* and *T. thiebautii*, respectively.[54] It would be most interesting to investigate if specialized metabolism plays a role in this niche differentiation. The discovery of a nondiazotrophic *Trichodesmium* species[55] provokes additional questions with respect to comparative metabolomics studies and the potential role of specialized metabolites in nitrogen fixation. Understanding how potential abiotic stressors regulate specialized metabolite and protein production will be key in determining the ecological role of the molecules we have discovered.[56]

## Conclusion

The impact of this study is that the incorporation of a large amount of high-resolution untargeted LC-MS/MS data offered the most elaborate temporal and spatial resolution for metabolite production in *Trichodesmium* known to the authors. We identified a new macrocylic polyketide-peptide (**1**), and we are well positioned to elucidate the communities, genetic architecture, and specialized metabolism that has allowed *Trichodesmium* species to attain prolific abundance in the oligotrophic open ocean. We focused on lipophilic metabolites in this work. This study still did not fully address the detection of saxitoxin and anatoxin found in previous work by research groups studying *Trichodesmium* blooms.[7,8] For instance, while we identified the widespread presence of the trichamide gene cluster in *T. thiebautii* H94 and several field samples, we did not detect it in any of our extracts, likely owing to its polarity. This could have been the case for the polar neurotoxins as well. Future studies will focus on the polar metabolites produced by *Trichodesmium* species. However, we are quite sure we would have captured microcystins in our extraction and fractionation approach as we used procedures similar to those of Briand and coworkers.[57] Further studies are still needed to fully understand if only certain species make these functionalized molecules, and what the physiological purpose of metabolites like trichothilone A (**1**) is, as it was found in extracts of *T. thiebautii* and *T. erythraeum*. For example, the presence of the euglenatides in multiple species of *Euglena* suggested an important biological role for these molecules.[58] Understanding specialized metabolite production in *Trichodesmium* species has application to areas as diverse as biological oceanography, chemical ecology, and drug discovery.

## Data Availability

All raw mass spectrometry files and .mzXML files can be found in the Center for Computational Mass Spectrometry MassIVE repository under number MSV0000930069. The raw NMR data and chemical shift information for trichothilone A (**1**) have been deposited at NP-MRD. 16S rRNA sequences have been deposited in NCBI GenBank (GoM2017 accession # OR661266; GoM2019-11 accession # OR665426). Information and data for the TriCoLim cruise[43] can be found at BCO-DMO https://www.bco-dmo.org/project/724451.

## Supporting information

Metabolite Annotations from Stations

## Acknowledgements

We profusely thank Dr. Simon J. Geist at Texas A&M, Corpus Christi for collecting the *Trichodesmium* bloom material in 2017. We gratefully acknowledge the crew of the Oregon II and NOAA for allowing us to volunteer on their annual ichthyoplankton survey in 2019 and for collecting samples in 2021, with special thanks to Andrew Millett and Glenn Zapfe. Acquisition of certain spectroscopic and spectrometric data in this publication was made possible by the use of equipment and services available through the RI-INBRE Centralized Research Core Facility at the University of Rhode Island, which is supported by the Institutional Development Award (IDeA) Network for Biomedical Research Excellence from the National Institute of General Medical Sciences of the National Institutes of Health under grant number P20GM103430. We also gratefully acknowledge support from NSF-2125191 to M.A.S and E.A.W., and the American Society of Pharmacognosy Starter Grant (M.J.B).

## Supplementary data

Supplementary data can be found with the online version of this article. It includes experimental methods, NMR and mass spectrometry data, computational data, phylogenetic analysis, and biosynthetic gene cluster information.

## References

[1] D. A. Hutchins, F.-X. Fu, E. A. Webb, N. Walworth, A. Tagliabue, Nature Geosci 2013, 6 (9), 790–795. 10.1038/ngeo1858.

[2] D. A. Hutchins, N. G. Walworth, E. A. Webb, M. A. Saito, D. Moran, M. R. McIlvin, J. Gale, F.-X. Fu, Nat. Commun. 2015, 6 (1), 8155. 10.1038/ncomms9155.

[3] T. G. Boatman, T. Lawson, R. J. Geider, PLoS ONE 2017, 12 (1), e0168796. 10.1371/journal.pone.0168796.

[4] S. P. Hawser, J. M. O’Neil, M. R. Roman, G. A. Codd, J. Appl. Phycol. 1992, 4 (1), 79–86. 10.1007/BF00003963.

[5] C. Guo, P. A. Tester, Nat. Toxins 1994, 2 (4), 222–227. 10.1002/nt.2620020411.

[6] S. Narayana, J. Chitra, S. R. Tapase, V. Thamke, P. Karthick, Ch. Ramesh, K. N. Murthy, M. Ramasamy, K. M. Kodam, R. Mohanraju, Harmful Algae 2014, 40, 34–39. 10.1016/j.hal.2014.10.003.

[7] A. M. Sacilotto Detoni, L. D. Fonseca Costa, L. A. Pacheco, J. S. Yunes, Toxicon 2016, 110, 51–55. 10.1016/j.toxicon.2015.12.003.

[8] S. Shunmugam, M. Gayathri, N. Prasannabalaji, N. Thajuddin, G. Muralitharan, Toxicon 2017, 135, 43–50. 10.1016/j.toxicon.2017.06.003.

[9] A. Ramos, A. Martel, G. Codd, E. Soler, J. Coca, A. Redondo, L. Morrison, J. Metcalf, A. Ojeda, S. Suárez, M. Petit, Mar. Ecol. Prog. Ser. 2005, 301, 303–305. 10.3354/meps301303.

[10] A. S. Kerbrat, Z. Amzil, R. Pawlowiez, S. Golubic, M. Sibat, H. T. Darius, M. Chinain, D. Laurent, Mar. Drugs 2011, 9 (4), 543–560. 10.3390/md9040543.

[11] S. Sudek, M. G. Haygood, D. T. A. Youssef, E. W. Schmidt, Appl. Environ. Microbiol. 2006, 72 (6), 4382–4387. 10.1128/AEM.00380-06.

[12] R. Endean, S. A. Monks, J. K. Griffith, L. E. Llewellyn, Toxicon 1993, 31 (9), 1155– 1165. 10.1016/0041-0101(93)90131-2.

[13] H. W. Paerl, B. M. Bebout, L. E. Prufert, J. Phycol. 1989, 25 (4), 773–784. 10.1111/j.0022-3646.1989.00773.x.

[14] P. J. A. Siddiqui, B. Bergman, E. J. Carpenter, Phycologia 1992, 31 (3–4), 326–337. 10.2216/i0031-8884-31-3-4-326.1.

[15] C. C. Sheridan, J. Plankton Res. 2002, 24 (9), 913–922. 10.1093/plankt/24.9.913.

[16] L. M. Momper, B. K. Reese, G. Carvalho, P. Lee, E. A. Webb, ISME J 2015, 9 (4), 882– 893. 10.1038/ismej.2014.186.

[17] M. Rouco, S. T. Haley, S. T. Dyhrman, Environ. Microbiol. 2016, 18, 5151–5160. 10.1111/1462-2920.13513.

[18] K. R. Frischkorn, M. Rouco, B. A. S. Van Mooy, S. T. Dyhrman, ISME J. 2017, 11, 2090–2101. 10.1038/ismej.2017.74.

[19] C. W. Via, E. Glukhov, S. Costa, P. V. Zimba, P. D. R. Moeller, W. H. Gerwick, M. J. Bertin, Front. Chem. 2018, 10.3389/fchem.2018.00316.

[20] M. J. Bertin, P. G. Wahome, P. V. Zimba, H. He, P. D. R. Moeller, Mar. Drugs 2017a, 15, 10.3390/md15010010.

[21] M. J. Bertin, J. Saurí, Y. Liu, C. W. Via, A. F. Roduit, R. T. Williamson. J. Org. Chem. 2018, 83, 13256–13266. 10.1021/acs.joc.8b02070.

[22] M. J. Bertin, A. F. Roduit, J. Sun, G. Alves, C. W. Via, M. A. Gonzalez, P. V. Zimba, P. D. R. Moeller, Mar. Drugs, 2017b, 15, 10.3390/md15070206.

[23] R. S. Belisle, C. W. Via, T. B. Schock, T. A. Villareal, P. V. Zimba, K. R. Beauchesne, P. D. R. Moeller, M. J. Bertin, Tetrahedron Lett. 2017, 58, 4066–4068. 10.1016/j.tetlet.2017.09.027.

[24] R. Teta, E. Irollo, G. Della Sala, G. Pirozzi, A. Mangoni, V. Costantino, Mar. Drugs 2013, 11 (11), 4451–4463. 10.3390/md11114451.

[25] G. Esposito, R. Teta, R. Miceli, L. Ceccarelli, G. Della Sala, R. Camerlingo, E. Irollo, A. Mangoni, G. Pirozzi, V. Costantino, Mar. Drugs 2015, 13 (1), 444–459. 10.3390/md13010444.

[26] M. J. Bertin, P. V. Zimba, H. He, P. D. R. Moeller, Tetrahedron Lett. 2016, 57, 5864– 5867. 10.1016/j.tetlet.2016.11.062.

[27] A. Caso, G. Esposito, G. Della Sala, J. R. Pawlik, R. Teta, A. Mangoni, V. Costantino, Mar. Drugs 2019, 17 (11), 618. 10.3390/md17110618.

[28] G. Esposito, G. Della Sala, R. Teta, A. Caso, M. Bourguet-Kondracki, J. R. Pawlik, A. Mangoni, V. Costantino, Eur. J. Org. Chem. 2016, 2016 (16), 2871–2875. 10.1002/ejoc.201600370.

[29] R. Teta, G. Della Sala, G. Esposito, C. W. Via, C. Mazzoccoli, C. Piccoli, M. J. Bertin, V. Costantino, A. Mangoni, Org. Chem. Front., 2019, 6, 1762–1774. 10.1039/C9QO00074G.

[30] K. M. McManus, R. D. Kirk, C. W. Via, J. S. Lotti, A. F. Roduit, R. Teta, S. Scarpato, A. Mangoni, M. J. Bertin, J. Nat. Prod. 2020, 83, 2664–2671. 10.1021/acs.jnatprod.0c00550.

[31] N. Matsumori, D. Kaneno, M. Murata, H. Nakamura, K. Tachibana, J. Org. Chem. 1999, 64 (3), 866–876. 10.1021/jo981810k.

[32] W. Koźmiński, D. Nanz, J. Magn. Reson. 2000, 142 (2), 294–299. 10.1006/jmre.1999.1939.

[33] P. Ciminiello, C. Dell’Aversano, E. Dello Iacovo, E. Fattorusso, M. Forino, L. Grauso, L. Tartaglione, Eur. J. Chem. 2012, 18 (52), 16836–16843. 10.1002/chem.201201357.

[34] Y. Kobayashi, J. Lee, K. Tezuka, Y. Kishi, Org. Lett. 1999, 1 (13), 2177–2180. 10.1021/ol9903786.

[35] J. Lee, Y. Kobayashi, K. Tezuka, Y. Kishi, Org. Lett. 1999, 1 (13), 2181–2184. 10.1021/ol990379y.

[36] L. Grauso, Y. Li, S. Scarpato, N. A. Cacciola, P. De Cicco, C. Zidorn, A. Mangoni, J. Nat. Prod. 2022, 85 (10), 2468–2473. 10.1021/acs.jnatprod.2c00796.

[37] S. Scarpato, R. Teta, P. De Cicco, F. Borrelli, J. R. Pawlik, V. Costantino, A. Mangoni, Mar. Drugs 2023, 21 (2), 58. 10.3390/md21020058.

[38] K. E. Gilbert, Pcmodel (version 10.0), Serena Software, Bloomington, IN, 2013.

[39] Gaussian 16, Revision C.01, Gaussian Inc., Wallingford CT, USA.

[40] N. Grimblat, M. M. Zanardi, A. M. Sarotti, J. Org. Chem. 2015, 80 (24), 12526–12534. 10.1021/acs.joc.5b02396.

[41] Y. Schmidt, K. Lehr, L. Colas, B. Breit, Chem. Eur. J. 2012,18, 7071–7081. 10.1002/chem.201103988.

[42] B. Bodo, W. Trowitzsch-Kienast, D. Schomburg, Tetrahedron Lett. 1986, 27 (7), 847–848. 10.1016/S0040-4039(00)84117-9.

[43] E. A. Webb, N. A. Held, Y. Zhao, E. D. Graham, A. E. Conover, J. Semones, M. D. Lee, Y. Feng, F. Fu, M. A. Saito, D. A. Hutchins, ISME Commun. 2023, 3 (1), 15. 10.1038/s43705-023-00214-y.

[44] K. Blin, S. Shaw, A. M. Kloosterman, Z. Charlop-Powers, G. P. van Weezel, M. H. Medema, T. Weber, Nucleic Acid Res. 2021, 49 (W1), W29–W35. 10.1093/nar/gkab335.

[45] Z. Chang, N. Sitachitta, J. V. Rossi, M. A. Roberts, P. M. Flatt, J. Jia, D. H. Sherman, W. H. Gerwick, J. Nat. Prod. 2004, 67 (8), 1356–1367. 10.1021/np0499261.

[46] J. J. Gehret, L. Gu, W. H. Gerwick, P. Wipf, D. H. Sherman, J. L. Smith, J. Biol. Chem. 2011, 286 (16), 14445–14454. 10.1074/jbc.M110.214635.

[47] G. M. Boratyn, A. A. Schäffer, R. Agarwala, S. F. Altschul, D. J. Lipman, T. L. Madden, Biol Direct 2012, 7, 12. 10.1186/1745-6150-7-12.

[48] D. J. Edwards, B. L. Marquez, L. M. Nogle, K. McPhail, D. E. Goeger, M. A. Roberts, W. H. Gerwick, Chem. Biol. 2004, 11 (6), 817–833. 10.1016/j.chembiol.2004.03.030.

[49] N. A. Moss, T. Leão, M. R. Rankin, T. M. McCullough, P. Qu, A. Korobeynikov, J. L. Smith, L. Gerwick, W. H. Gerwick, ACS Chem. Biol. 2018, 13 (12), 3385–3395. 10.1021/acschembio.8b00910.

[50] B. Bergman, G. Sandh, S. Lin, J. Larsson, E. J. Carpenter, FEMS Microbiol. Rev. 2013, 37, 286–302. 10.1111/j.1574-6976.2012.00352.x.

[51] A. M. Hynes, E. A. Webb, S. C. Doney, J. B. Waterbury, J. Phycol. 2012, 48 (1), 196–210. 10.1111/j.1529-8817.2011.01096.x.

[52] N. A. Moss, G. Seiler, T. F. Leão, G. Castro-Falcón, L. Gerwick, C. C. Hughes, W. H. Gerwick, Angew. Chem. Int. Ed. 2019, 58 (27), 9027–9031. 10.1002/anie.201902571.

[53] N. A. Held, K. M. Sutherland, E. A. Webb, M. R. McIlvin, N. R. Cohen, A. J. Devaux, D. A. Hutchins, J. B. Waterbury, C. M. Hansel, M. A. Saito, ISME Commun. 2021, 1, 35. 10.1038/s43705-021-00034-y.

[54] K. A. Confesor, C. R. Selden, K. E. Powell, L. A. Donahue, T. Mellett, S. Caprara, A. N. Knapp, K. N. Buck, P. D. Chappell, Front. Mar. Sci. 2022, 9, 10.3389/fmars.2022.821655.

[55] T. O. Delmont, Proc. Natl. Acad. Sci. U. S. A. 2021, 118 (46), e2112355118. 10.1073/pnas.2112355118.

[56] N. A. Held, E. A. Webb, M. M. McIlvin, D. A. Hutchins, N. R. Cohen, D. M. Moran, K. Kunde, M. C. Lohan, C. Mahaffey, E. M. S. Woodward, M. A. Saito, Biogeosciences 2020, 17, 2537–2551. 10.5194/bg-17-2537-2020.

[57] E. Briand, M. Bormans, M. Gugger, P. C. Dorrestein, W. H. Gerwick, Environ. Microbiol. 2016, 18 (2), 384–400. 10.1111/1462-2920.12904.

[58] M. Aldholmi, R. Ahmad, D. Carretero-Molina, I. Pérez-Victoria, J. Martín, F. Reyes, O. Genilloud, L. Gourbeyre, T. Gefflaut, H. Carlsson, A. Maklakov, E. O’Neill, R. A. Field, B. Wilkinson, M. O’Connell, A. Ganesan, Angew. Chem. Int. Ed. 2022, 61, e202203175. 10.1002/anie.202203175.

