## Supporting Information for "Spatial and Temporal Resolution of Cyanobacterial Bloom Chemistry Reveals an Open-Ocean *Trichodesmium thiebautii* as a Talented Producer of Specialized Metabolites"

[b] Dr. L. Grauso
Dipartimento di Agraria, Università degli Studi di Napoli Federico II, via Universita 100, 80055 Portici Napoli, Italy

[c] Prof. E. A. Webb, Dr. N. A. Held
Marine and Environmental Biology, Department of Biological Sciences, University of Southern California, Los Angeles, CA 90089, United States

[d] Dr. M. A. Saito
Department of Marine Chemistry and Geochemistry, Woods Hole Oceanographic Institution, Woods Hole, MA 02543, United States

[e] Dr. S. Scarpato, Prof. A. Mangoni
Dipartimento di Farmacia, Università degli Studi di Napoli Federico II, via Domenico Montesano 49, 80131 Napoli, Italy

[f] Dr. P. V. Zimba
Rice Rivers Center, Virginia Commonwealth University, Richmond, VA 23284, United States

[g] Dr. P. D. R. Moeller
Harmful Algal Bloom Monitoring and Reference Branch, Stressor Detection and Impacts Division, National Ocean Service/NOAA, Hollings Marine Laboratory, 331 Fort Johnson Road, Charleston, SC 29412, United States

[h] Dr. M. J. Bertin
Department of Chemistry, Case Western Reserve University, Cleveland, OH 44106, United States

+These authors contributed equally.

**Table of Contents**

**1. Experimental Section……………………………………………………………………….3**

**1.1 General experimental procedures……………………………………………...3**

**1.2 Collection of cyanobacteria, genetic analysis, and extraction procedures………….3**

**1.3 Isolation of 1…………………………………………………………………………..4**

**1.4 Application of Kishi's method……………………………………………………….5**

**1.5 Preparation and Analysis of MTPA esters…………………………………………6**

**1.6 Methanolysis of 1……………………………………………………………………..6**

**1.7 Computational details………………………………………………………………..7**

**1.8 LC-MS/MS-based molecular networking……………………………………………..8**

**1.9 *Trichodesmium* spp extracts: mass spectrometry assay using LC-MS/MS and the GNPS library…………………………………………………………………9**

**1.10 Cytotoxicity assays………………..………………………...………………...10**

**1.11 Genomic analysis…………………………………………………………………..10**

**2. Supplementary tables……………………………………………………………………..12**

**3. Supplementary figures……………………………………………………………………18**

**4. References………………………………………………………………………………44**

### 1. Experimental Section

**1.1 General experimental procedures.** Optical rotation was measured using a Jasco P-2000 polarimeter. UV spectra were measured using a Beckman Coulter DU-800 spectrophotometer. IR spectra were obtained using a Thermo Scientific Nicolet 380 FT-IR spectrometer. NMR spectra were recorded on a Varian 500 MHz NMR instrument and the trichothilone A methyl ester (**2**) was analyzed using a Bruker Ascend 600 MHz NMR instrument with $CD_3OD$ (referenced to residual $CH_3OH$ at $\delta_H$ 4.78 and $\delta_H$ 3.31 and $\delta_C$ 49.2), $(CD_3)_2SO$ (referenced to residual DMSO at $\delta_H$ 2.50 and $\delta_C$ 39.5) or $CDCl_3$ (referenced to residual $CHCl_3$ at $\delta_H$ 7.26 and $\delta_C$ 77.2). HRESIMS analysis was performed using an AB SCIEX TripleTOF 4600 mass spectrometer with Analyst TF software. Additional high-resolution LC-MS/MS data was collected on a Thermo LTQ Orbitrap XL coupled to a Dionex Ultimate 3000 HPLC. Low resolution LC-MS/MS data was collected on a Thermo LTQ XL coupled to a Dionex Ultimate 3000 HPLC. Semi-preparative HPLC was carried out using a Dionex UltiMate 3000 HPLC system equipped with a micro vacuum degasser, an autosampler and a diode-array detector.

**1.2 Collection of cyanobacteria, genetic analysis, and extraction procedures.** *Trichodesmium* biomass was collected from the Gulf of Mexico near North Padre Island, TX in 2014. As described previously, the dominant organism was identified as *Trichodesmium* sp., and the material was shipped to our laboratory.[1,2] The biomass (14 g dry weight) was repeatedly extracted with a 2:1 mixture of $CH_2Cl_2$:$CH_3OH$ and the crude extract was concentrated to an oil under reduced pressure. Collections were also made from the GoM in 2017, 2019 and 2021. Select samples were preserved in RNAlater and transferred to the laboratory. DNA isolation, PCR amplification of 16S rRNA genes, and phylogenetic analysis followed the procedures of McManus and coworkers exactly.[3] 16S rRNA sequences have been deposited in NCBI GenBank (GoM2017 accession #

OR661266; GoM2019-11 accession # OR665426). Additionally, samples were acquired from the TriCoLim 2018 expedition,[4] and from culture collections (Table S1). The biomass at each site was extracted in 2:1 $CH_2Cl_2$:$CH_3OH$, followed by a 1:1 $CH_2Cl_2$:$CH_3OH$, and 100% $CH_3OH$. The resulting crude extracts were separately concentrated to oils under reduced pressure. Small aliquots of the resulting crude extracts were passed over a 100 mg C18 SPE column to prepare for LC-MS/MS analysis. The extract from a bulk collection of biomass (GOM2019-6) was reconstituted in hexanes and fractionated over silica gel using the same method as the North Padre Island 2014 collection,[2] creating nine fractions (A-I). Fraction H (25% $CH_3OH$ in EtOAc) was further fractioned over a 2-gram C18 SPE column into two fractions, eluting with 100% $CH_3OH$ and 100% EtOAc. The 100% $CH_3OH$ fraction (27.8 mg), was subjected to reversed phase HPLC using a 5µm Kinetex C18 (250 x 10 mm) semi preparative column utilizing a gradient and a flow rate at 3.0 ml/min of $H_2O$ with 0.05% formic acid and $CH_3CN$ with 0.05% formic acid. The gradient elution was as follows: 50% $CH_3CN$ for 3 min, 50% to 100% $CH_3CN$ for 19 min, 100% for 10 min and then a return to initial conditions. Collections were made every 5 min starting at 10 min, the resulting collection between 25 and 30 min was combined with the PI2014 HPLC fraction (100% $CH_3CN$) from which **1** was isolated (described below).

**1.3 Isolation of 1.** The oily residue from the PI2014 extract was reconstituted in hexanes and fractionated over silica gel using a stepwise gradient from hexanes to ethyl acetate to methanol generating nine fractions total (A-I).[2] Fractions G (100% EtOAc) and H (25% $CH_3OH$ in EtOAc) were combined based on similarities in $^1$H NMR spectra and MS profiles. The combined fractions were further fractionated using a 2 g C18 SPE column and four additional fractions were generated eluting with 50% $CH_3CN$ in $H_2O$, 100% $CH_3CN$, 100% $CH_3OH$ and 100% EtOAc. The 100% $CH_3CN$ fraction (143.2 mg) was combined with the GOM2019-6 fraction described above and subjected to reversed phase HPLC using a Kinetex 5 μm C18 column (250 x 10 mm) utilizing an isocratic gradient of 65% $CH_3CN$ in $H_2O$ modified with 0.05% formic acid and a flow rate of 3.0 mL/min. This resulted in the isolation of 8.9 mg of **1** ($t_R$, 27.00 min). The raw NMR data and chemical shift information for trichothilone A (**1**) have been deposited at NP-MRD.

*Trichothilone A* (**1**): pale yellow oil; $[\alpha]^{23}_{D}$ +8.5 (*c* 0.10, $CH_3OH$); UV ($CH_3OH$) $\lambda_{max}$ (log $\varepsilon$) 202 (3.5), 244 (3.0) nm; IR (ZnSe) $\nu_{max}$ 3391 (br), 2925, 1730, 1605, 1376, 1192 $cm^{-1}$, $^{1}H$ NMR (500 MHz, DMSO-$d_6$) and $^{13}C$ NMR (125 MHz, DMSO-$d_6$), see Table 1, HRMS (ESI) *m/z* calcd for $C_{30}H_{53}N_2O_4S$: 537.3721 $[M + H]^+$; found 537.3725 (error 0.74 ppm).

**1.4 Application of Kishi's method.** We utilized Kishi's method to provide support for the relative configuration analysis of the contiguous propionate segment in **1**,[5,6] spanning C-14 to C-17. Chemdraw 12 was used to predict chemical shifts for an acyclic simplified model compound of trichothilone A (**1**) to provide corrected chemical shifts for the comparative analysis to the database compounds 1a-1h.

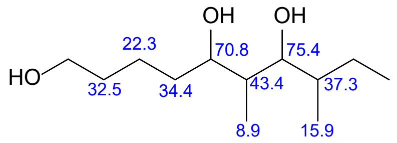


reference compound (**1a-h**)

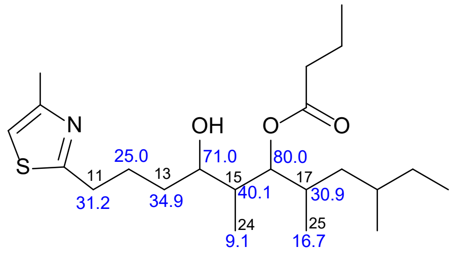


simplified acyclic analogue of trichothilone A

Using these corrected chemical shifts, the differences between predicted and experimental shifts were determined and delta $^{13}C$ NMR values were calculated (Table S2). The best configuration match was to stereoisomer A (ααββ). Additionally, examining mean of absolute differences (MAE) again matched stereoisomer A as the best match.

**1.5 Preparation and Analysis of MTPA esters.** 3 mg of **1** was dissolved in dry $CDCl_3$ and separated into two equal portions in 4 mL vials. Dry pyridine (10 µL) and (*S*)-(+)-α-methoxy-α-(trifluoromethyl)phenylacetyl chloride (15 µL) were added to the first vial. The vial was capped and the reaction mixture was stirred for 24 h. The identical procedure was repeated with an equal amount of **1** and (*R*)-(–)-α-methoxy-α-(trifluoromethyl)phenylacetyl chloride. These reactions yielded the C-14 *R* ester from (*S*)-MTPA-Cl and the C-14 *S* ester from (*R*)-MTPA-Cl. Each reaction mixture was separated between $CH_2Cl_2$ and water, and the organic phase was evaporated and subjected to reversed phase HPLC using a Kinetex C18 column (250 x 10 mm, 5 µm) and an isocratic method using 100% $CH_3CN$. The purified esters were analyzed by $^1$H NMR, COSY, and TOCSY to firmly establish $^1$H NMR chemical shifts of derivatives. For partial $^1$H NMR chemical shift information of derivatives see Figure S18.

**1.6 Methanolysis of 1.** 5.3 mg of **1** were transferred to a 4 mL vial and stirred with 2 mL of 2 N HCl in 92% $CH_3OH$ for 16 h at 65 °C. Next, the reaction mixture was dried under a stream of $N_2$ and the presence of the expected *m/z* 659 of the methyl ester of **1** was checked by LC-MS/MS. The reaction mixture was subjected to reversed phase HPLC using a Kinetex C18 column (250 x 10 mm, 5 µm) and an isocratic method using 65% $CH_3CN$ in $H_2O$ modified with 0.5% formic acid. The trichothilone A methyl ester (**2**) eluted at 11.5 min (0.6 mg) and was characterized by high resolution mass spectrometry and $^1$H NMR. COSY and multiplicity-edited HSQC spectra were used to determine the proton chemical shifts of H-18a and H-18b in the 1,3-dimethyl system in **2**.

*Trichothilone A methyl ester* (**2**): colorless oil; $^1$H NMR (600 MHz, DMSO-$d_6$) $\delta$ 7.06 (s, 1H; H-9), 4.03 (m, 1H; H-14), 3.84 (m, 1H; H-16), 3.58 (s, 3H; H-22), 3.21 (m, 1H; H-21a) 3.17

(m, 1H; H-21b), 2.94 (t, *J*=8 Hz, 1H; H-11a), 2.90/2.76 (s, 3H; H-28), 2.60 (m, 2H; H-7), 2.54 (m, 1H; H-11b), 2.46 (m, 1H; H-2); 2.27 (m, 2H; H-29), 1.82 (m, 1H; H-12a), 1.68 (m, 1H; H-12b) 1.60 (m, 1H; H17), 1.58 (m, 2H; H-6), 1.50 (m, 1H; H-4), 1.43 (ovlp, 1H; H-15), 1.42 (ovlp, 2H; H-3), 1.42 (ovlp, 1H; 19) 1.40 (ovlp, 1H; H-20a), 1.33 (ovlp, 1H; H-18a), 1.31 (ovlp, 1H; H-5a), 1.30 (ovlp, 1H; H-20b); 1.23 (m, 2H; H-13), 1.15 (m, 1H; H-5b), 1.04 (d, *J*=6.7 Hz, 3H; H-23), 0.95 (m, 3H; H-30) 0.90 (m, 1H; H-18b), 0.85 (m, 3H; H-24), 0.81 (d, J=6.1 Hz, 3H; H-27), 0.72 (m, 3H; H-26), 0.68 (d, J=6.9 Hz, 3H; H-25); HRMS (ESI) *m/z* calcd for $C_{31}H_{57}N_2O_5S$: 569.3983 $[M + H]^+$; found 569.3984 (error 0.18 ppm).

**1.7 Computational details.** Conformational search for the four diastereomers at C-2 and C-4 of model compound **1m** (*RR*-**1m**, *RS*-**1m**, *SR*-**1m**, *SS*-**1m**) was performed using the program Pcmodel v. 10[7] and the GMMX algorithm in the MMFF94 force field. The search generated, respectively, 69, 50, 66, and 35 conformers for the four diastereomers within 4 kcal/mol from the lowest-energy conformer. These conformers were used as starting structure for density functional theory (DFT) calculations, performed using the program Gaussian 16.[8] The geometry of each conformer was optimized at the B3LYP/6-31G(d) level of theory. The population of each optimized conformer was calculated using the Boltzmann distribution law at 298 K and the energy calculated at the B3LYP/6-31G(d,p) level of theory with the SMD model for the solvent, DMSO. The resulting 16, 13, 14, and 11 conformers with population >1% for, respectively, *RR*-**1m**, *RS*-**1m**, *SR*-**1m**, and *SS*-**1m**, were used for the subsequent NMR calculations. The lowest-energy conformer of each stereoisomer is depicted in Figure S15. The Cartesian coordinates of all conformers used to calculate NMR parameters are provided as Supporting Information in the separate text file Compounds_1m_coord.txt, and the isotropic shieldings for each conformer in the separate Excel file Isotropic_shieldings.xlsx. NMR calculations were performed at the mPW1PW91/6-

311+G(d,p) level of theory, with the PCM model for the solvent. The isotropic shielding calculated for each nucleus were averaged over the conformers according to their respective populations, and average isotropic shielding were converted into chemical shifts using the scaling factors proposed by Pierens[9] or used directly for application of the DP4+ method.[10] Chemical shifts of C-16, C-17, C-18, C-25, H-17, $H_3$-18, and $H_3$-25 were not included in the comparison between predicted and experimental chemical shifts, because the predicted chemical shifts of these nuclei were remarkably affected by the shorter side chain of model compounds **1m** compared to the natural product **1**. Average isotropic shieldings and predicted chemical shifts for all the **1m** diastereomers are reported in Table S4, and the detailed results of DP4+ calculations are reported in Table S5.

**1.8 LC-MS/MS-based molecular networking.** The resulting extracts described above were analyzed on a liquid chromatography system, a Dionex Ultimate 3000 HPLC, coupled to high resolution electrospray mass spectrometer, a Thermo LTQ Orbitrap XL system. The LC portion included an in-line degasser, binary pump and refrigerated auto sampler. The column oven was maintained at room temperature, and a 5 μm Kinetex C18 column (50 x 2.1 mm) was used with a flow of 200 μL/min with $H_2O$ (0.1% formic acid) and $CH_3OH$. The gradient elution was as follows: 45% $CH_3OH$ for 1 min, 45%-80% $CH_3OH$ over 30 min, 100% $CH_3OH$ for 9 min. All the mass spectrometry data were collected in positive ion detection mode. The MS spray voltage was 4.8 kV with a capillary temperature of 280 °C. After each full-scan MS spectrum, the five most intense ions were selected for fragmentation in five subsequent MS/MS scans. The CID isolation width was 3.0 and the normalized collision energy was set to 35.0 units with activation time 30 ms. The raw data files were transformed to .mzXML files using the publicly available MSConvert. Data were uploaded and networked using the online platform at GNPS (https://gnps.ucsd.edu/).[10] The parent mass tolerance and MS/MS fragment mass tolerance were both set to the default 0.02 Da for high resolution data. The networks cosine score for edge connection was set to 0.6 with at least 3 matched peaks required. The spectra were searched in the built in GNPS library of spectra, where matched nodes need to have at least 6 matched peaks and meet the minimum cosine score of 0.7. A background blank sample was filtered against the network to remove those ions. The resulting networks were downloaded and further visualized in Cytoscape 3.9.1. Metabolites were annotated by comparing retention time, HRMS values, and product ion profiles to a pure compound library of metabolites isolated from the 2014 investigation and those metabolites previously characterized from *Smenospongia aurea*. All raw mass spectrometry files and .mzXML files can be found in the Center for Computational Mass Spectrometry MassIVE repository under number MSV0000930069.

**1.9 *Trichodesmium* spp extracts: mass spectrometry assay using LC-MS/MS and the GNPS library.** The MS/MS data for all metabolites characterized in our previous work (with the exceptions of trichophycin I, trichotoxin B, trichophycin D, trichophycin E, and **1**) were added to the GNPS library (see Table S6 for library ID numbers).[10] Extracts from GOM 2021 (stations 2, 3, and 4) and the TriCoLim expedition (Stations 3, 4, 6, 8, 13 15, 16, 17, and 20) were analyzed by LC-MS/MS. Raw data were collected on a Dionex Ultimate 3000 HPLC system coupled to a Thermo Scientific LTQ XL mass spectrometer. The LC portion included a binary pump, refrigerated auto sampler and a column oven kept at 30 ºC. A 2.6 µm Kinetex C18 column (150 x 4.6 mm) was used for separations with a flow rate set at 400 µL/min. A gradient method consisting of $H_2O$ with 0.1% formic acid and $CH_3CN$ with 0.1% formic acid was used. The gradient method was as follows: 50% $CH_3CN$ was held for 5 min, followed by a 15 min gradient of 50% $CH_3CN$ to 100% $CH_3CN$, which was held for 10 min followed by a return to initial conditions from 31-38 min. The MS spray voltage was 3.5 kV with a capillary temperature of 325 °C. For the MS/MS component, the CID isolation width was 1.0 and the collision energy was 35.0 eV. Once acquisitions were complete, .raw data files were exported and transformed into .mzXML format using MSConvert. Files were uploaded into the GNPS platform for molecular networking. The parent mass tolerance and MS/MS fragment mass tolerance were set to the default 2.0 Da and 0.5 Da, respectively for low resolution data. The networks cosine score for edge connection was set to 0.6 with at least 2 matched peaks required. The spectra were searched in the built in GNPS library of spectra, where matched nodes need to have at least 2 matched peaks and meet the minimum cosine score of 0.6. Library hits were annotated for each sample and verified by retention time comparisons to standards and MS/MS fragmentation comparisons. This method was also used to detect **1** from IMS101 cells and media extracts.

**1.10 Cytotoxicity assays.** Assays were carried out as previously described.[2] Briefly, Neuro-2A cells were added to assay plates in 100 µl of Eagle's Minimum Essential Media (EMEM) supplemented with 10% FBS at a density of 5,000 cells/well. Cells were incubated overnight (37 °C, 5% $CO_2$) and examined microscopically to confirm confluence and adherence. Compound **1** was dissolved in DMSO (1% v/v) and added to the cells in the range of 100 to 0.1 µM in order to generate $EC_{50}$ curves against both cell lines. Four technical replicates were prepared for each concentration and each assay was performed in triplicate. Doxorubicin was used as a positive control and DMSO (1% v/v) was used as a negative control. Assays were resolved after 72 h by the addition of MTT reagent to test and control wells, a 4 h incubation period, aspiration of media, and addition of DMSO to dissolve formazan crystals. Absorbance was recorded at 540 nm using a SpectraMax plate reader and $EC_{50}$ curves were generated using Graphpad Prism software.

**1.11 Genomic analysis.** The *Trichodesmium erythraeum* IMS101 assembled genome sequence was retrieved from the European Nucleotide Archive (accession #CP000393). The *T. thiebautii*

H94 genome and a group of field sample metagenome-assembled genome (MAG) assemblies were retrieved from NCBI's SRA (BioProject PRJNA828267). Assemblies were downloaded as FASTA files and uploaded to the antiSMASH platform where putative biosynthetic gene clusters were identified and annotated. Webb and coworkers isolated DNA from the samples and sequenced and assembly MAGs following the R/V Atlantis TriCoLim cruise,[4] and these samples have some colocation with the archived samples from the cruise that we analyzed via mass spectrometry during this work. Information and data for the TriCoLim cruise can be found at BCO-DMO https://www.bco-dmo.org/project/724451.

**Table S1**. Sample location sites in this study.

| Sample ID | Species ID | GPS Coordinates | Collection Notes[b] |
|---|---|---|---|
| PI2014 | *T. thiebautii* | ND[a] | Bulk biomass |
| GoM2017 | *T. thiebautii* | 27.00ºN; 92.00ºW | Bulk biomass |
| GoM2019-2 | ND | 26.00ºN; 93.59ºW | Picked colonies (mixed morphology) |
| GoM2019-3 | ND | 26.18ºN; 94.39ºW | Picked colonies (mixed morphology) |
| GoM2019-4 | ND | 26.01ºN; 94.59ºW | Picked colonies (mixed morphology) |
| GoM2019-6 | ND | 28.00ºN; 93.00ºW | Bulk biomass |
| GoM2019-11 | *T. thiebautii* | 29.00ºN; 88.00ºW | Picked colonies (mixed morphology) |
| GoM2021-2 | ND | 26.00ºN; 93.59ºW | Bulk biomass |
| GoM2021-3 | ND | 26.00ºN; 93.29ºW | Bulk biomass |
| GoM2021-4 | ND | 28.00ºN; 90.00ºW | Bulk biomass |
| Tricolim St 3 | ND | 9.893ºN; 22.241ºW | Bulk biomass |
| Tricolim St 4 | ND | 6.011ºN; 21.593ºW | Picked colonies (mixed morphology) |
| Tricolim St 6 | ND | 22.19ºN; 35.53ºW | Bulk biomass |
| Tricolim St 8 | ND | 3.724ºS; 22.0116ºW | Bulk biomass |
| Tricolim St 13 | ND | 0.976ºS; 30.843ºW | GFF filter |
| Tricolim St 15 | ND | 5.239ºN; 44.958ºW | Picked colonies (only puffs) |
| Tricolim St 16 | ND | 7.012ºN; 48.958ºW | Picked colonies (mixed morphology) |
| Tricolim St 17 | ND | 10.760ºN; 55.849ºW | Picked colonies (mixed morphology) |
| Tricolim St 20 | ND | 16.862ºN; 65.036ºW | Picked colonies (mixed morphology) |
| ST8 | *T. erythraeum* | Culture from USCTCC | |
| IMS101 | *T. erythraeum* | Culture from Bigelow Labs | |

[a]not determined; [b]colony morphology: puff or tuft or mixed

**Table S2.** Determination of the relative configuration of the two contiguous propionate units in **1** by comparison to NMR database[33] – analysis in three NMR solvents ($CD_3OD$, DMSO-*d*$_6$, $CDCl_3$). Differences in $^{13}C$ NMR signals were determined and mean absolute values of differences determined the best fit to the database, which was configuration A: α, α, β, β (cf. Figure S14).

Experimental chemical shifts of apralide A were corrected as suggested in the Kishi's paper before comparison with database data, using chemical shift predicted by ChemDraw 12.0 and the compounds below:

| | CD3OD | | | | DMSO | | | | CDCl3 | | |
|---|---|---|---|---|---|---|---|---|---|---|---|
| | Exp | Correctior | Corrected | | Exp | Correctior | Corrected | | Exp | Correctior | Corrected |
| 11 | 32.2 | -1.3 | 33.5 | | 31.2 | -1.3 | 32.5 | | 31.8 | -1.3 | 33.1 |
| 12 | 26.6 | 2.7 | 23.9 | | 26.0 | 2.7 | 23.3 | | 25.8 | 2.7 | 23.1 |
| 13 | 33.4 | 0.5 | 32.9 | | 32.5 | 0.5 | 32.0 | | 34.5 | 0.5 | 34.0 |
| 14 | 71.4 | -0.2 | 71.6 | | 68.9 | -0.2 | 69.1 | | 69.3 | -0.2 | 69.5 |
| 15 | 40.7 | -3.3 | 44.0 | | 38.8 | -3.3 | 42.1 | | 38.6 | -3.3 | 41.9 |
| 16 | 78.0 | 4.6 | 73.4 | | 76.7 | 4.6 | 72.1 | | 77.2 | 4.6 | 72.6 |
| 17 | 31.9 | -6.4 | 38.3 | | 30.5 | -6.4 | 36.9 | | 30.7 | -6.4 | 37.1 |
| 24 | 9.5 | 0.2 | 9.3 | | 9.4 | 0.2 | 9.2 | | 8.9 | 0.2 | 8.7 |
| 25 | 12.6 | 0.8 | 11.8 | | 12.3 | 0.8 | 11.5 | | 12.4 | 0.8 | 11.6 |

| | CD3OD | 13C | | | | | | | | | | | | | | | | | | | | | | |
|---|---|---|---|---|---|---|---|---|---|---|---|---|---|---|---|---|---|---|---|---|---|---|---|---|
| | A, ααββ | | | B, αααα | | | C, ααβα | | | D, αααβ | | | E, βαββ | | | F, βααα | | | G, βαβα | | | H, βααβ | | |
| Position | Exp | Database | Delta | Exp | Database | Delta | Exp | Database | Delta | Exp | Database | Delta | Exp | Database | Delta | Exp | Database | Delta | Exp | Database | Delta | Exp | Database | Delta |
| 11 | 33.5 | 33.7 | -0.2 | 33.5 | 33.7 | -0.2 | 33.5 | 33.7 | -0.2 | 33.5 | 33.7 | -0.2 | 33.5 | 33.7 | -0.2 | 33.5 | 33.7 | -0.2 | 33.5 | 33.7 | -0.2 | 33.5 | 33.7 | -0.2 |
| 12 | 23.9 | 23.9 | 0.0 | 23.9 | 23.5 | 0.4 | 23.9 | 23.7 | 0.2 | 23.9 | 23.4 | 0.5 | 23.9 | 23.2 | 0.7 | 23.9 | 23.5 | 0.4 | 23.9 | 23.2 | 0.7 | 23.9 | 23.4 | 0.5 |
| 13 | 32.9 | 35.3 | -2.4 | 32.9 | 35.9 | -3.0 | 32.9 | 35.5 | -2.6 | 32.9 | 35.7 | -2.8 | 32.9 | 33.0 | -0.1 | 32.9 | 33.4 | -0.5 | 32.9 | 33.4 | -0.5 | 32.9 | 35.7 | -2.8 |
| 14 | 71.6 | 72.6 | -1.0 | 71.6 | 75.2 | -3.6 | 71.6 | 72.6 | -1.0 | 71.6 | 76.7 | -5.1 | 71.6 | 75.0 | -3.4 | 71.6 | 75.0 | -3.4 | 71.6 | 75.0 | -3.4 | 71.6 | 75.5 | -3.9 |
| 15 | 44.0 | 41.1 | 2.9 | 44.0 | 40.5 | 3.5 | 44.0 | 39.7 | 4.3 | 44.0 | 39.9 | 4.1 | 44.0 | 42.7 | 1.3 | 44.0 | 42.6 | 1.4 | 44.0 | 42.6 | 1.4 | 44.0 | 40.5 | 3.5 |
| 16 | 73.4 | 76.4 | -3.0 | 73.4 | 79.0 | -5.6 | 73.4 | 80.1 | -6.7 | 73.4 | 80.2 | -6.8 | 73.4 | 77.6 | -4.2 | 73.4 | 80.6 | -7.2 | 73.4 | 80.6 | -7.2 | 73.4 | 75.7 | -2.3 |
| 17 | 38.3 | 38.0 | 0.3 | 38.3 | 38.2 | 0.1 | 38.3 | 38.6 | -0.3 | 38.3 | 38.8 | -0.5 | 38.3 | 38.1 | 0.2 | 38.3 | 38.4 | -0.1 | 38.3 | 38.8 | -0.5 | 38.3 | 38.8 | -0.5 |
| 24 | 9.3 | 10.5 | -1.2 | 9.3 | 7.8 | 1.5 | 9.3 | 11.0 | -1.7 | 9.3 | 6.6 | 2.7 | 9.3 | 11.7 | -2.4 | 9.3 | 12.2 | -2.9 | 9.3 | 10.0 | -0.7 | 9.3 | 10.1 | -0.8 |
| 25 | 11.8 | 12.6 | -0.8 | 11.8 | 14.6 | -2.8 | 11.8 | 16.6 | -4.8 | 11.8 | 15.5 | -3.7 | 11.8 | 12.1 | -0.3 | 11.8 | 17.2 | -5.4 | 11.8 | 15.4 | -3.6 | 11.8 | 15.4 | -3.6 |
| Sum of absolute errors | | | 11.8 | | | 20.7 | | | 21.8 | | | 26.4 | | | 12.8 | | | 21.5 | | | 18.2 | | | 18.1 |
| Mean absolute errors | | | 1.3 | | | 2.3 | | | 2.4 | | | 2.9 | | | 1.4 | | | 2.4 | | | 2.0 | | | 2.0 |

| | DMSO | 13C | | | | | | | | | | | | | | | | | | | | | | |
|---|---|---|---|---|---|---|---|---|---|---|---|---|---|---|---|---|---|---|---|---|---|---|---|---|
| | A, ααββ | | | B, αααα | | | C, ααβα | | | D, αααβ | | | E, βαββ | | | F, βααα | | | G, βαβα | | | H, βααβ | | |
| Position | Exp | Database | Delta | Exp | Database | Delta | Exp | Database | Delta | Exp | Database | Delta | Exp | Database | Delta | Exp | Database | Delta | Exp | Database | Delta | Exp | Database | Delta |
| 11 | 32.5 | 32.7 | -0.2 | 32.5 | 32.8 | -0.3 | 32.5 | 32.8 | -0.3 | 32.5 | 33.0 | -0.5 | 32.5 | 32.8 | -0.3 | 32.5 | 32.8 | -0.3 | 32.5 | 32.7 | -0.2 | 32.5 | 32.8 | -0.3 |
| 12 | 23.3 | 22.7 | 0.6 | 23.3 | 22.3 | 1.0 | 23.3 | 22.5 | 0.8 | 23.3 | 22.1 | 1.2 | 23.3 | 22.1 | 1.2 | 23.3 | 22.2 | 1.1 | 23.3 | 22.3 | 1.0 | 23.3 | 22.1 | 1.2 |
| 13 | 32.0 | 34.2 | -2.2 | 32.0 | 34.8 | -2.8 | 32.0 | 34.4 | -2.4 | 32.0 | 34.6 | -2.6 | 32.0 | 34.6 | -2.6 | 32.0 | 33.3 | -1.3 | 32.0 | 31.8 | 0.2 | 32.0 | 34.2 | -2.2 |
| 14 | 69.1 | 69.6 | -0.5 | 69.1 | 72.1 | -3.0 | 69.1 | 69.7 | -0.6 | 69.1 | 73.8 | -4.7 | 69.1 | 71.7 | -2.6 | 69.1 | 71.9 | -2.8 | 69.1 | 71.6 | -2.5 | 69.1 | 72.5 | -3.4 |
| 15 | 42.1 | 39.8 | 2.3 | 42.1 | 39.1 | 3.0 | 42.1 | 38.5 | 3.6 | 42.1 | 38.5 | 3.6 | 42.1 | 41.6 | 0.5 | 42.1 | 39.9 | 2.2 | 42.1 | 41.4 | 0.7 | 42.1 | 39.2 | 2.9 |
| 16 | 72.1 | 73.2 | -1.1 | 72.1 | 75.6 | -3.5 | 72.1 | 77.0 | -4.9 | 72.1 | 77.1 | -5.0 | 72.1 | 74.5 | -2.4 | 72.1 | 72.9 | -0.8 | 72.1 | 77.4 | -5.3 | 72.1 | 72.9 | -0.8 |
| 17 | 36.9 | 36.2 | 0.7 | 36.9 | 36.5 | 0.4 | 36.9 | 36.8 | 0.1 | 36.9 | 37.0 | -0.1 | 36.9 | 36.4 | 0.5 | 36.9 | 36.9 | 0.0 | 36.9 | 36.7 | 0.2 | 36.9 | 37.2 | -0.3 |
| 24 | 9.2 | 10.0 | -0.8 | 9.2 | 8.0 | 1.2 | 9.2 | 10.6 | -1.4 | 9.2 | 6.8 | 2.4 | 9.2 | 11.2 | -2.0 | 9.2 | 10.3 | -1.1 | 9.2 | 11.5 | -2.3 | 9.2 | 9.8 | -0.6 |
| 25 | 11.5 | 12.3 | -0.8 | 11.5 | 14.1 | -2.6 | 11.5 | 16.5 | -5.0 | 11.5 | 15.3 | -3.8 | 11.5 | 12.0 | -0.5 | 11.5 | 15.1 | -3.6 | 11.5 | 16.9 | -5.4 | 11.5 | 15.1 | -3.6 |
| Sum of absolute errors | | | 9.2 | | | 17.8 | | | 19.1 | | | 23.9 | | | 12.6 | | | 13.2 | | | 17.8 | | | 15.3 |
| Mean absolute errors | | | 1.0 | | | 2.0 | | | 2.1 | | | 2.7 | | | 1.4 | | | 1.5 | | | 2.0 | | | 1.7 |

| | CDCl3 | 13C | | | | | | | | | | | | | | | | | | | | | | |
|---|---|---|---|---|---|---|---|---|---|---|---|---|---|---|---|---|---|---|---|---|---|---|---|---|
| | A, ααββ | | | B, αααα | | | C, ααβα | | | D, αααβ | | | E, βαββ | | | F, βααα | | | G, βαβα | | | H, βααβ | | |
| Position | Exp | Database | Delta | Exp | Database | Delta | Exp | Database | Delta | Exp | Database | Delta | Exp | Database | Delta | Exp | Database | Delta | Exp | Database | Delta | Exp | Database | Delta |
| 11 | 33.1 | 32.6 | 0.5 | 33.1 | 32.6 | 0.5 | 33.1 | 32.5 | 0.6 | 33.1 | 32.5 | 0.6 | 33.1 | 32.5 | 0.6 | 33.1 | 32.5 | 0.6 | 33.1 | 32.5 | 0.6 | 33.1 | 32.5 | 0.6 |
| 12 | 23.1 | 22.8 | 0.3 | 23.1 | 22.5 | 0.6 | 23.1 | 22.6 | 0.5 | 23.1 | 22.5 | 0.6 | 23.1 | 21.2 | 1.9 | 23.1 | 22.6 | 0.5 | 23.1 | 21.3 | 1.8 | 23.1 | 22.6 | 0.5 |
| 13 | 34.0 | 33.0 | 1.0 | 34.0 | 35.1 | -1.1 | 34.0 | 33.9 | 0.1 | 34.0 | 35.1 | -1.1 | 34.0 | 34.2 | -0.2 | 34.0 | 35.1 | -1.1 | 34.0 | 34.3 | -0.3 | 34.0 | 35.2 | -1.2 |
| 14 | 69.5 | 73.5 | -4.0 | 69.5 | 77.0 | -7.5 | 69.5 | 72.3 | -2.8 | 69.5 | 77.4 | -7.9 | 69.5 | 76.7 | -7.2 | 69.5 | 76.5 | -7.0 | 69.5 | 76.5 | -7.0 | 69.5 | 76.6 | -7.1 |
| 15 | 41.9 | 39.0 | 2.9 | 41.9 | 37.9 | 4.0 | 41.9 | 37.8 | 4.1 | 41.9 | 37.8 | 4.1 | 41.9 | 40.9 | 1.0 | 41.9 | 38.2 | 3.7 | 41.9 | 40.8 | 1.1 | 41.9 | 38.1 | 3.8 |
| 16 | 72.6 | 77.6 | -5.0 | 72.6 | 81.5 | -8.9 | 72.6 | 80.4 | -7.8 | 72.6 | 81.4 | -8.8 | 72.6 | 79.3 | -6.7 | 72.6 | 75.3 | -2.7 | 72.6 | 82.0 | -9.4 | 72.6 | 74.9 | -2.3 |
| 17 | 37.1 | 37.0 | 0.1 | 37.1 | 37.6 | -0.5 | 37.1 | 37.5 | -0.4 | 37.1 | 37.7 | -0.6 | 37.1 | 36.8 | 0.3 | 37.1 | 37.3 | -0.2 | 37.1 | 37.0 | 0.1 | 37.1 | 37.6 | -0.5 |
| 24 | 8.7 | 11.7 | -3.0 | 8.7 | 4.7 | 4.0 | 8.7 | 11.6 | -2.9 | 8.7 | 4.2 | 4.5 | 8.7 | 13.0 | -4.3 | 8.7 | 10.8 | -2.1 | 8.7 | 13.4 | -4.7 | 8.7 | 10.5 | -1.8 |
| 25 | 11.6 | 12.8 | -1.2 | 11.6 | 15.1 | -3.5 | 11.6 | 15.8 | -4.2 | 11.6 | 14.9 | -3.3 | 11.6 | 11.6 | 0.0 | 11.6 | 15.3 | -3.7 | 11.6 | 16.7 | -5.1 | 11.6 | 14.8 | -3.2 |
| Sum of absolute errors | | | 18.0 | | | 30.6 | | | 23.4 | | | 31.5 | | | 22.2 | | | 21.6 | | | 30.1 | | | 21.0 |
| Mean absolute errors | | | 2.0 | | | 3.4 | | | 2.6 | | | 3.5 | | | 2.5 | | | 2.4 | | | 3.3 | | | 2.3 |

**Table S3**. Metabolites found in GoM 2021 and TriCoLim samples

| Sample ID | Metabolites |
|---|---|
| GoM2021-2 | putative smenamide A/B analog, trichophycin F, trichothiazole A |
| GoM2021-3 | smenamides C/D, trichophycin F |
| GoM2021-4 | putative smenamide A/B analog, smenamides C/D, trichophycin F, trichothiazole A |
| Tricolim St 3 | smenamides A/B, trichophycin F, trichothiazole A |
| Tricolim St 4 | ND[a] |
| Tricolim St 6 | trichophycin F |
| Tricolim St 8 | trichothiazole A |
| Tricolim St 13 | ND |
| Tricolim St 15 | ND |
| Tricolim St 16 | trichothiazole A |
| Tricolim St 17 | trichophycin F, trichothiazole A |
| Tricolim St 20 | ND |

[a]none detected

**Table S4**. Average isotropic shielding constants and predicted chemical shifts of the four stereoisomers of model compound **1m** calculated at the mPW1PW91/6-311+G(d,p)/PCM(DMSO) level, compared to experimental chemical shifts of compound **1**.

| | Average shieldings[a] | | | | Predicted chemical shifts[b] | | | | Experimental chemical shifts |
|---|---|---|---|---|---|---|---|---|---|
| Position | *RR*-**1m** | *RS*-**1m** | *SR*-**1m** | *SS*-**1m** | *RR*-**1m** | *RS*-**1m** | *SR*-**1m** | *SS*-**1m** | |
| C-1 | -0.8 | -1.9 | -2.1 | -3.1 | 178.2 | 179.2 | 179.5 | 180.4 | 175.5 |
| C-2 | 143.7 | 144.2 | 146.2 | 146.1 | 40.5 | 40.0 | 38.2 | 38.3 | 35.5 |
| C-3 | 141.8 | 143.5 | 143.4 | 140.9 | 42.4 | 40.7 | 40.8 | 43.2 | 40.0 |
| C-4 | 152.9 | 154.4 | 154.0 | 153.5 | 31.8 | 30.4 | 30.7 | 31.2 | 29.1 |
| C-5 | 149.9 | 151.0 | 150.5 | 149.7 | 34.6 | 33.6 | 34.1 | 34.8 | 34.9 |
| C-6 | 157.8 | 157.6 | 157.8 | 156.9 | 27.1 | 27.3 | 27.1 | 28.0 | 25.0 |
| C-7 | 153.9 | 154.1 | 153.7 | 154.8 | 30.8 | 30.6 | 31.0 | 30.0 | 29.9 |
| C-8 | 24.8 | 24.7 | 25.1 | 24.8 | 153.8 | 153.9 | 153.5 | 153.8 | 156.0 |
| C-11 | 152.3 | 152.5 | 151.9 | 152.4 | 32.3 | 32.1 | 32.8 | 32.2 | 31.2 |
| C-12 | 159.2 | 157.5 | 156.5 | 157.7 | 25.8 | 27.4 | 28.3 | 27.2 | 26.0 |
| C-13 | 154.0 | 155.3 | 154.6 | 154.0 | 30.7 | 29.5 | 30.1 | 30.7 | 32.5 |
| C-14 | 116.6 | 117.5 | 115.9 | 115.8 | 66.3 | 65.5 | 67.1 | 67.1 | 68.9 |
| C-15 | 146.1 | 145.1 | 145.4 | 143.8 | 38.3 | 39.2 | 38.9 | 40.4 | 38.8 |
| C-22 | 168.2 | 169.9 | 169.9 | 167.3 | 17.2 | 15.6 | 15.5 | 18.1 | 16.4 |
| C-23 | 169.4 | 167.6 | 167.3 | 170.0 | 16.0 | 17.8 | 18.1 | 15.5 | 19.9 |
| C-24 | 178.2 | 178.4 | 178.1 | 178.1 | 7.7 | 7.5 | 7.8 | 7.8 | 9.4 |
| | | | | **$^{13}$C RMSD** | **2.32** | **2.24** | **1.98** | **2.46** | |
| H-2 | 29.06 | 29.06 | 28.99 | 29.00 | 2.51 | 2.52 | 2.58 | 2.57 | 2.44 |
| H-3a | 30.28 | 30.32 | 30.41 | 30.29 | 1.36 | 1.32 | 1.24 | 1.36 | 1.32 |
| H-3b | 30.37 | 30.42 | 30.45 | 30.50 | 1.27 | 1.23 | 1.20 | 1.16 | 1.16 |
| H-4 | 30.27 | 29.94 | 30.17 | 29.98 | 1.37 | 1.68 | 1.47 | 1.64 | 1.42 |
| H-5a | 30.18 | 30.17 | 30.31 | 30.15 | 1.45 | 1.47 | 1.34 | 1.48 | 1.06 |
| H-5b | 30.69 | 30.67 | 30.68 | 30.63 | 0.98 | 0.99 | 0.99 | 1.03 | 1.06 |
| H-6a | 29.82 | 29.89 | 29.92 | 29.90 | 1.80 | 1.73 | 1.71 | 1.72 | 1.70 |
| H-6b | 30.19 | 30.10 | 30.05 | 30.21 | 1.45 | 1.54 | 1.58 | 1.43 | 1.59 |
| H-7a | 28.97 | 28.90 | 28.84 | 28.97 | 2.60 | 2.67 | 2.73 | 2.60 | 2.73 |
| H-7b | 29.04 | 29.12 | 29.19 | 29.01 | 2.53 | 2.46 | 2.39 | 2.56 | 2.49 |
| H-11a | 28.70 | 28.78 | 28.61 | 28.68 | 2.86 | 2.78 | 2.94 | 2.87 | 2.96 |
| H-11b | 28.78 | 28.79 | 28.89 | 28.76 | 2.78 | 2.77 | 2.68 | 2.80 | 2.77 |
| H-12a | 29.80 | 29.82 | 29.90 | 29.82 | 1.81 | 1.80 | 1.72 | 1.80 | 1.75 |
| H-12b | 30.14 | 29.93 | 29.92 | 29.82 | 1.50 | 1.69 | 1.70 | 1.80 | 1.75 |
| H-13a | 30.12 | 30.21 | 30.14 | 30.18 | 1.51 | 1.43 | 1.49 | 1.45 | 1.40 |
| H-13b | 30.38 | 30.56 | 30.56 | 30.65 | 1.27 | 1.10 | 1.09 | 1.01 | 1.13 |
| H-14 | 28.36 | 28.38 | 28.52 | 28.32 | 3.18 | 3.16 | 3.02 | 3.21 | 3.18 |
| H-15 | 30.11 | 30.19 | 30.20 | 30.20 | 1.53 | 1.45 | 1.44 | 1.44 | 1.51 |
| H-16 | 27.02 | 27.05 | 27.05 | 27.05 | 4.44 | 4.41 | 4.42 | 4.42 | 4.81 |
| H3-22 | 30.57 | 30.58 | 30.65 | 30.65 | 1.09 | 1.08 | 1.02 | 1.02 | 1.01 |
| H3-23 | 30.78 | 30.88 | 30.89 | 30.84 | 0.89 | 0.80 | 0.79 | 0.83 | 0.81 |
| H3-24 | 30.93 | 30.95 | 30.95 | 30.94 | 0.75 | 0.73 | 0.73 | 0.74 | 0.71 |
| | | | | **$^{1}$H RMSD** | **0.150** | **0.145** | **0.122** | **0.149** | |

[a] This is the set of shielding used for DP4+ analysis. Results are reported in Table S4.
[b] Chemical shifts were calculated according to ref. 52, using the equations $\delta = (186.2534 - \text{shielding})/1.0496$ for $^{13}$C chemical shift and $\delta = (31.7217 - \text{shielding})/1.058$ for $^{1}$H chemical shifts.

**Table S5.** Detailed DP4+ analysis of predicted chemical shifts for the four stereoisomers of model compound **1m** compared to experimental chemical shifts of compound **1**.

| Functional | Solvent? | Basis Set? | Type of Data |
|---|---|---|---|
| mPW1PW91 | PCM | 6-311+G(d,p) | Shielding Tensors |

| | *RR*-**1m** | *RS*-**1m** | *SR*-**1m** | *SS*-**1m** | Isomer 5 | Isomer 6 |
|---|---|---|---|---|---|---|
| sDP4+ (H data) | 0.95% | 2.42% | 96.42% | 0.21% | - | - |
| sDP4+ (C data) | 5.07% | 9.79% | 83.11% | 2.03% | - | - |
| sDP4+ (all data) | 0.06% | 0.29% | 99.64% | 0.01% | - | - |
| uDP4+ (H data) | 0.29% | 2.67% | 95.84% | 1.20% | - | - |
| uDP4+ (C data) | 4.44% | 8.06% | 84.74% | 2.76% | - | - |
| uDP4+ (all data) | 0.02% | 0.26% | 99.68% | 0.04% | - | - |
| DP4+ (H data) | 0.00% | 0.07% | 99.92% | 0.00% | - | - |
| DP4+ (C data) | 0.32% | 1.10% | 98.50% | 0.08% | - | - |
| DP4+ (all data) | 0.00% | 0.00% | 100.00% | 0.00% | - | - |

**Table S6.** GNPS Library ID numbers for *Trichodesmium* specialized metabolites.

| Compound Name | GNPS Library ID |
|---|---|
| trichotoxin A | CCMSLIB00011427499 |
| trichothiazole A | CCMSLIB00006678013 |
| trichophycin G | CCMSLIB00011427498 |
| conulothiazole A | CCMSLIB00011427497 |
| isotrichophycin C/trichophycin C | CCMSLIB00011427491 |
| isoconulothiazole B/conulothiazole B | CCMSLIB00006678009 |
| tricholide A | CCMSLIB00006710031 |
| smenolactone D | CCMSLIB00011427490 |
| trichophycin B/smenolactone C | CCMSLIB00006678007 |
| trichophycin B/smenolactone C ($Na^+$ adduct) | CCMSLIB00011427548 |
| tricholide B | CCMSLIB00006710032 |
| conulothiazole C | CCMSLIB00006678010 |
| trichophycin F | CCMSLIB00006678011 |
| trichophycin H | CCMSLIB00011427492 |
| smenothiazole B | CCMSLIB00011427493 |
| smenamide C/D | CCMSLIB00004752853 |
| trichophycin A | CCMSLIB00006678012 |
| smenothiazole A | CCMSLIB00011427494 |
| smenamide E | CCMSLIB00011427495 |
| smenamide A/B | CCMSLIB00004752852 |
| smenamide F | CCMSLIB00011427496 |
| trichothilone A | CCMSLIB00012176070 |
| unnarmicin D | CCMSLIB00006678014 |

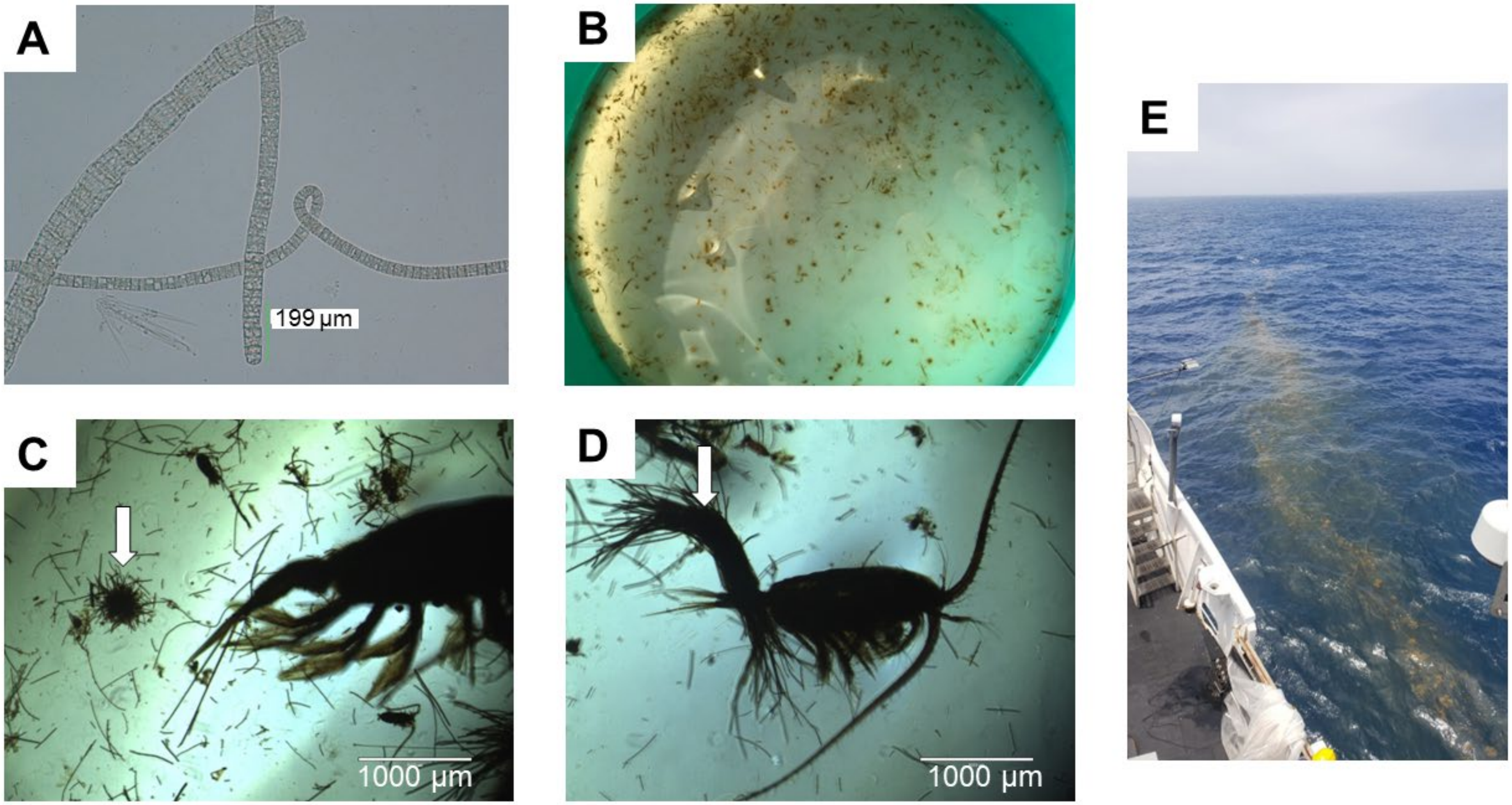


Figure S1. (A) Microscopy of *Trichodesmium* field collection from 2017. (B) Puff and tuft colonies of *Trichodesmium* in collection sieve at GoM2019-11. (C) Micrograph of puff colony (arrow). (D) Micrograph of tuft colony. (E) Surface accumulation from GoM2019-6.

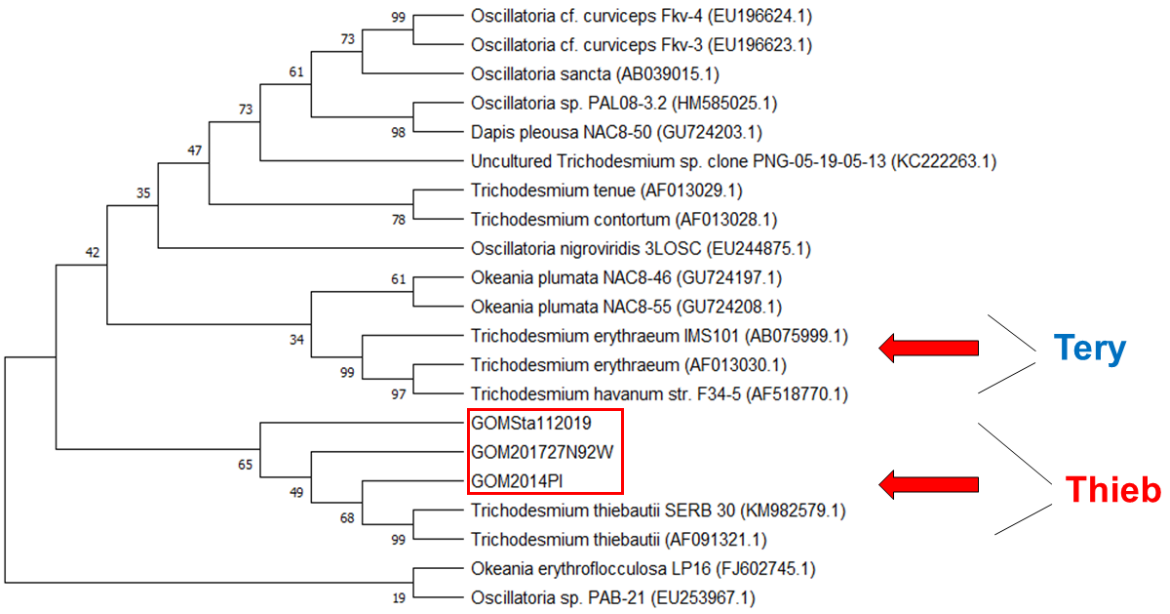


Figure S2. 16S rRNA phylogenetic tree aligning *Trichodesmium* species from this study (red box - PI2014, GoM 2017, and GoM2019-11). All samples clustered with *T. thiebautii*. The tree was created using the Maximum Likelihood method and the Tamura-Nei model. The bootstrap consensus tree is inferred from 1000 replicates and the percentage of replicate trees in which the associated taxa clustered together in the bootstrap test are shown next to branches. Analysis was conducted in MEGA. Genbank accession numbers of sequences are noted in parentheses. Red arrow show clusters for *T. erythraeum* (Tery) and *T. thiebautii* (Thieb).

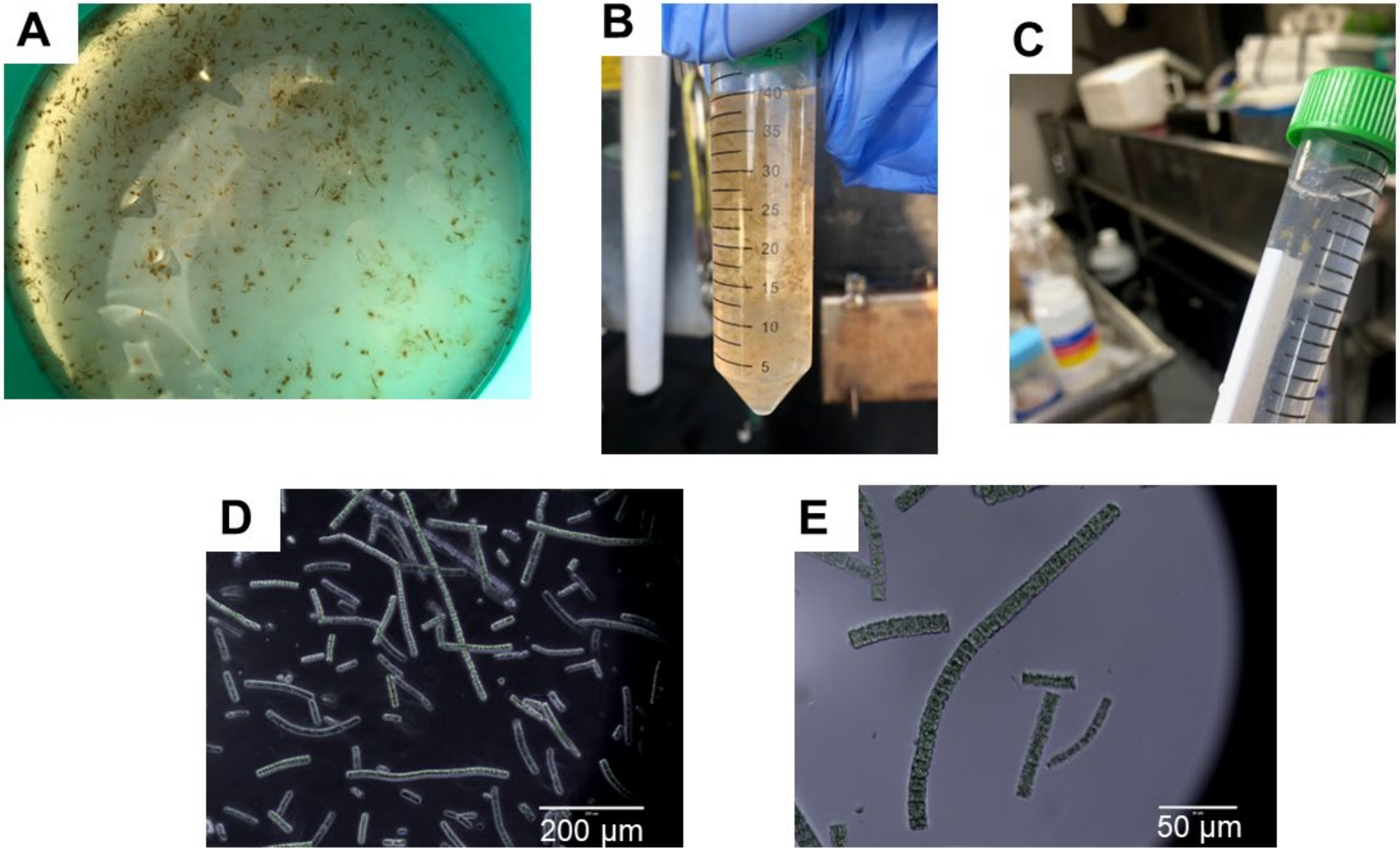


Figure S3. The process of "colony picking" from GoM2019-11. (A) Colonies collected in a tow net were washed in a sieve. (B) Colonies in the sieve were transferred to a 50 mL falcon tube. (C) Colonies were transferred with a sterile loop one-by-one to a 15 mL falcon tube. (D, E) Micrographs of filaments after the processed was completed.

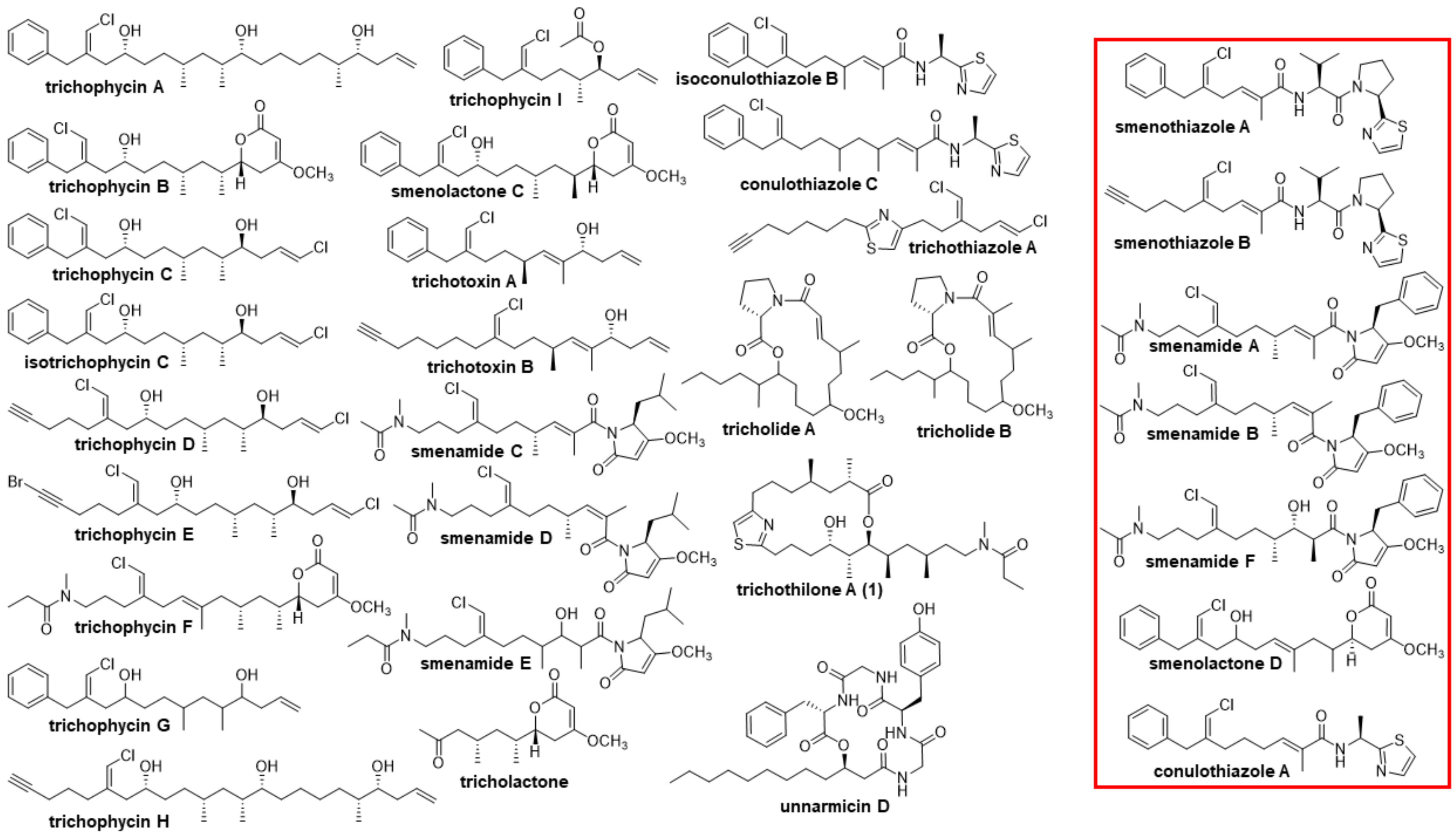


Figure S4. “Compound library”: the 31 compounds that were isolated or detected from the PI2014 collection. The molecules in the red box designate those molecules isolated from the sponge *Smenospongia aurea* but also detected in *T. thiebautii*.

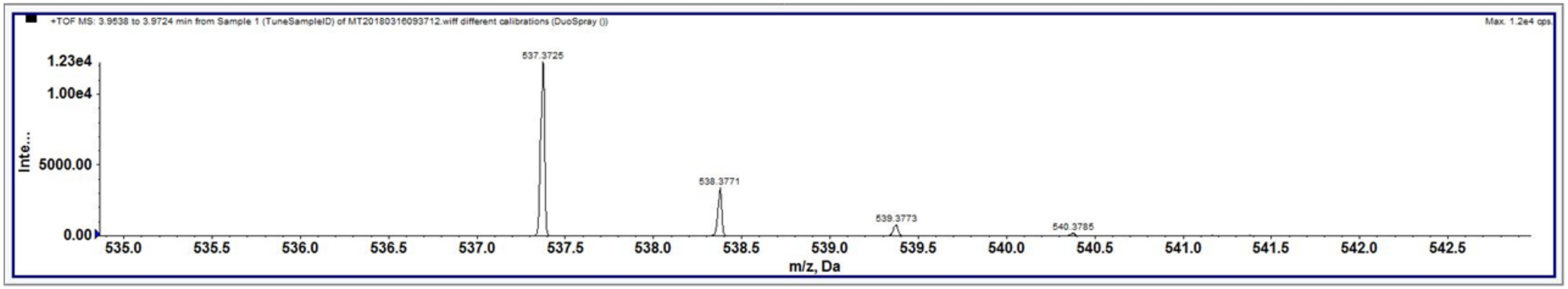


Figure 5. HRESIMS of (**1**) with *m/z* 537.3725 $[M+H]^+$ found.

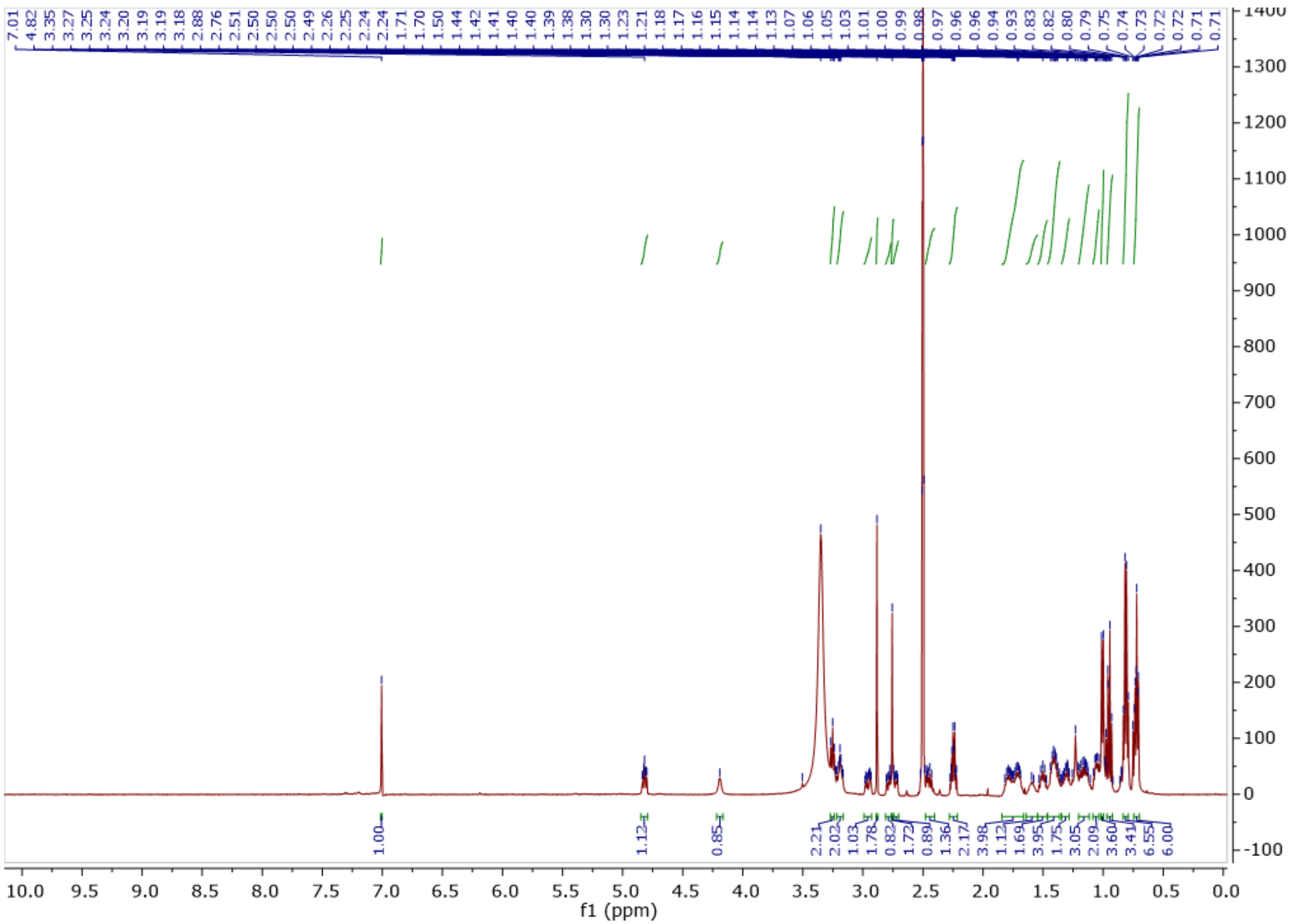

Figure S6. $^{1}$H NMR of trichothilone (**1**) (500 MHz, DMSO-$d_6$).

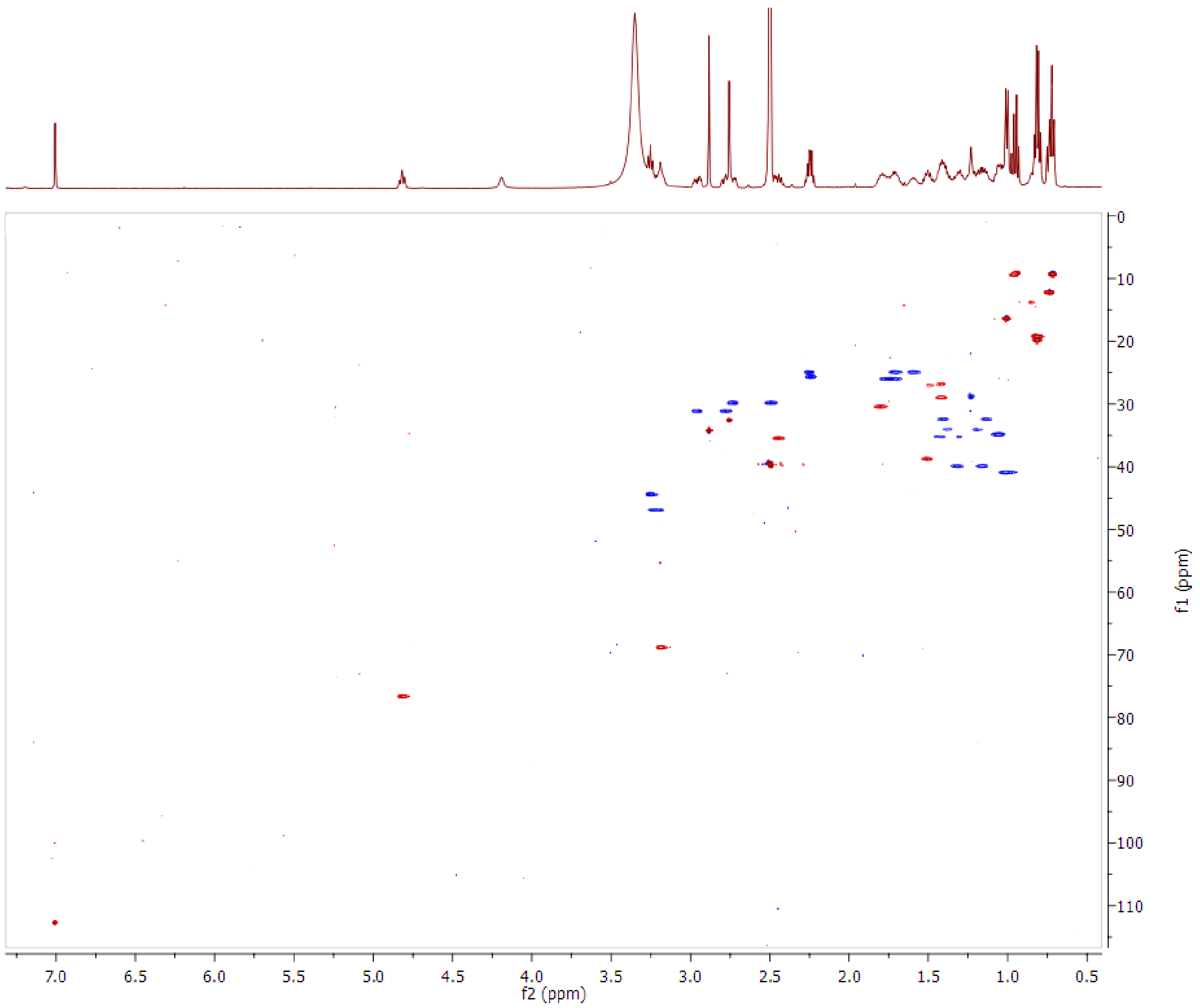


Figure S7. Multiplicity-edited HSQC of **1**.

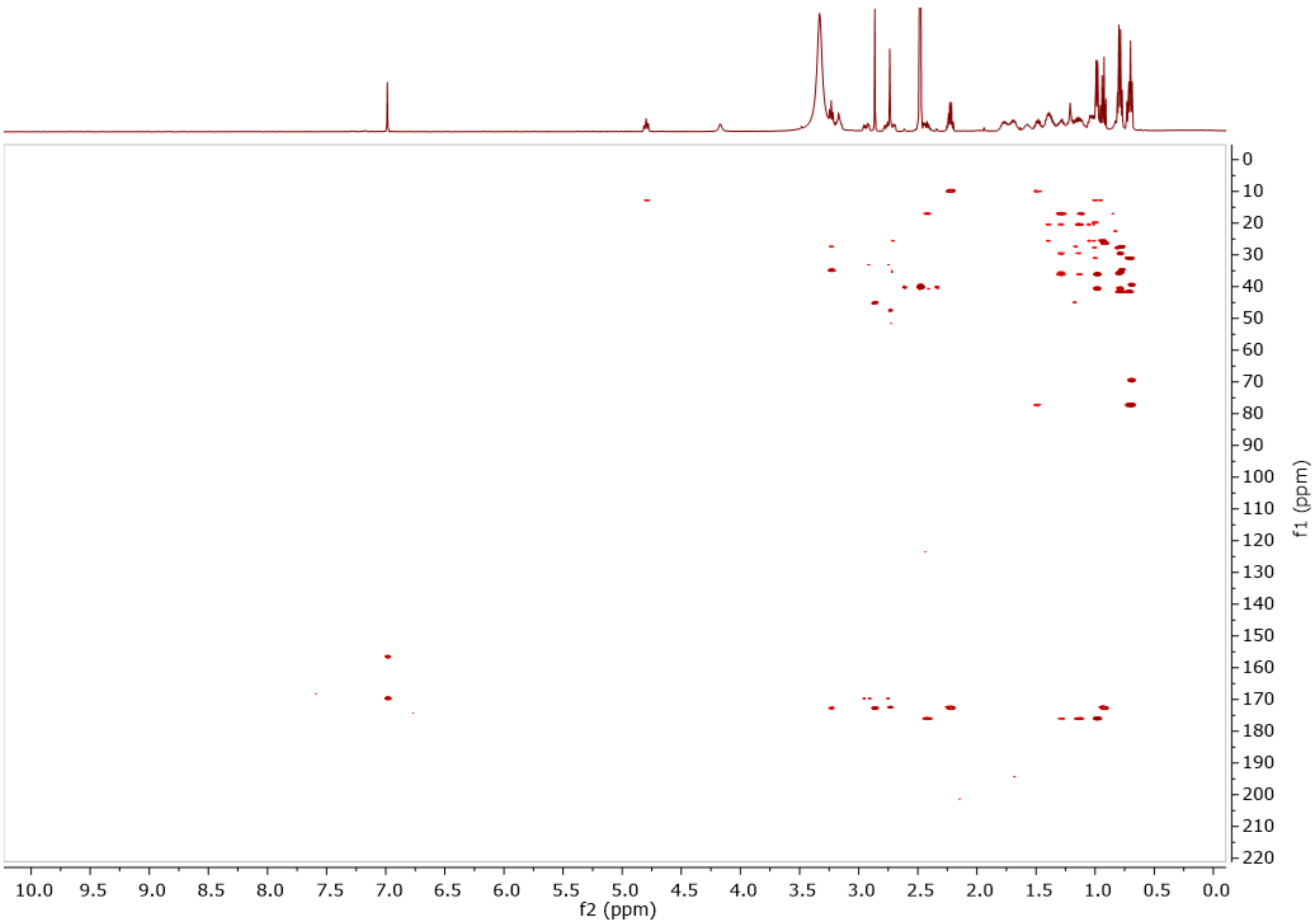


Figure S8. HMBC of **1**.

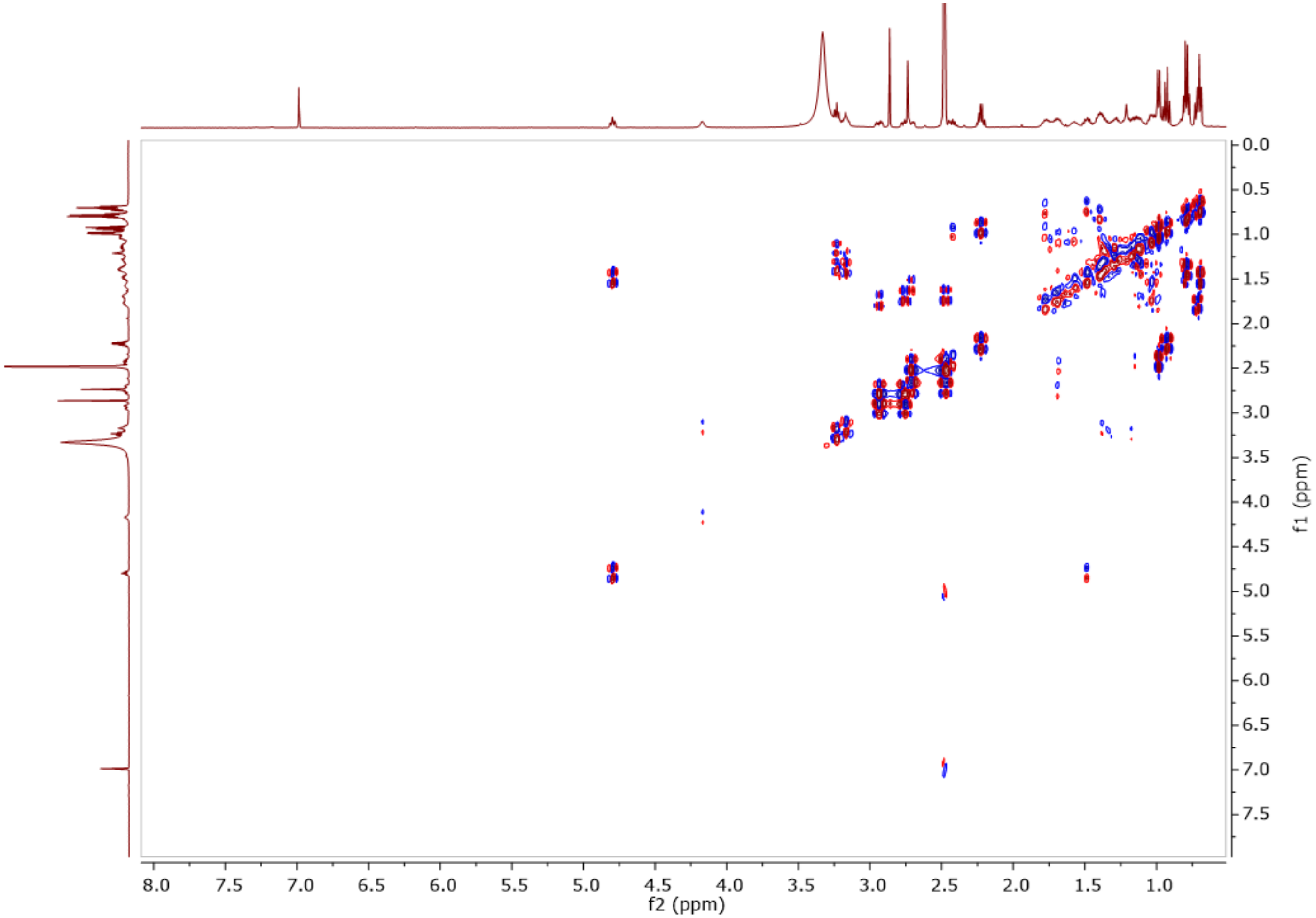


Figure S9. DQF-COSY of **1**.

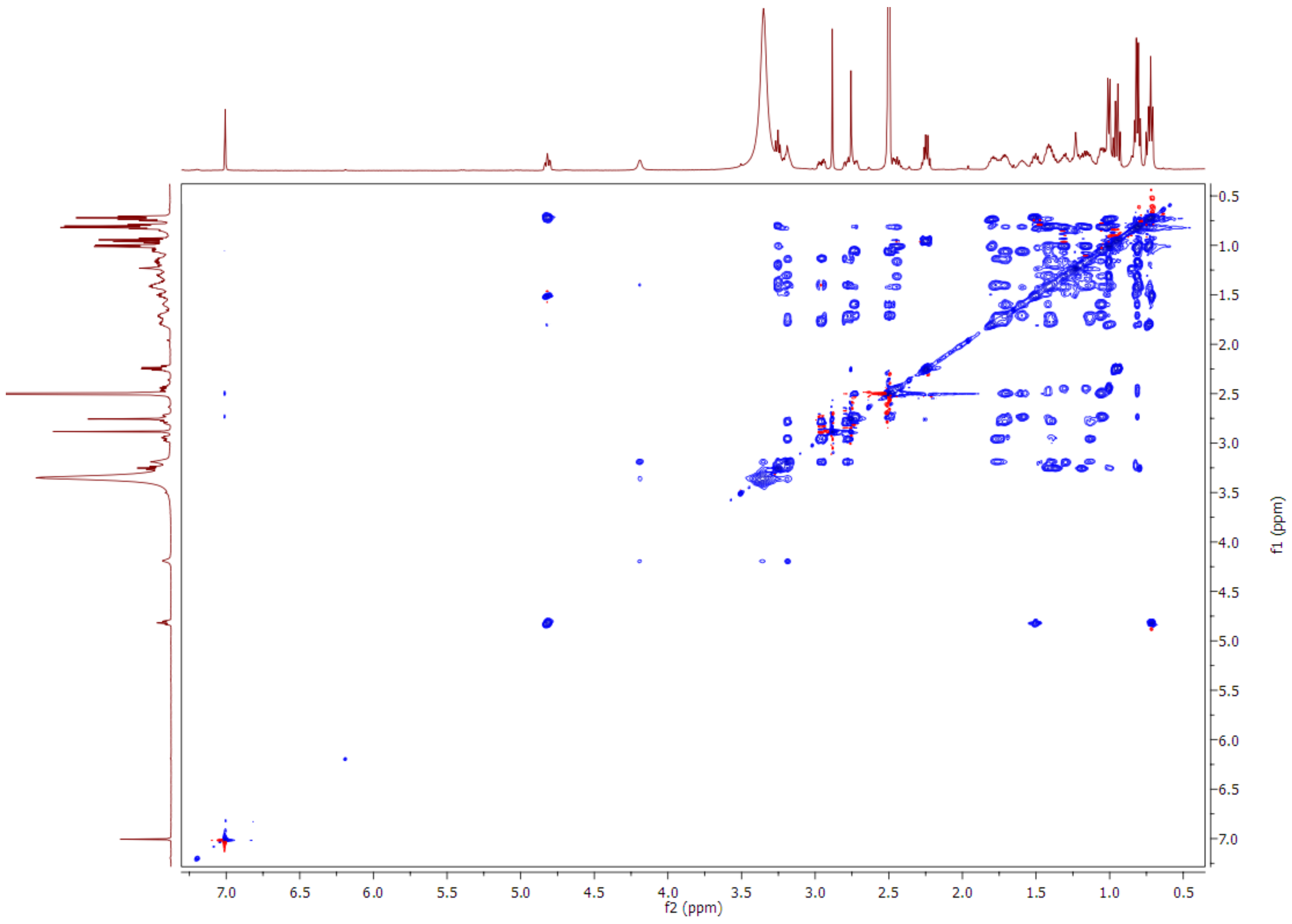


Figure S10. TOCSY of **1**.

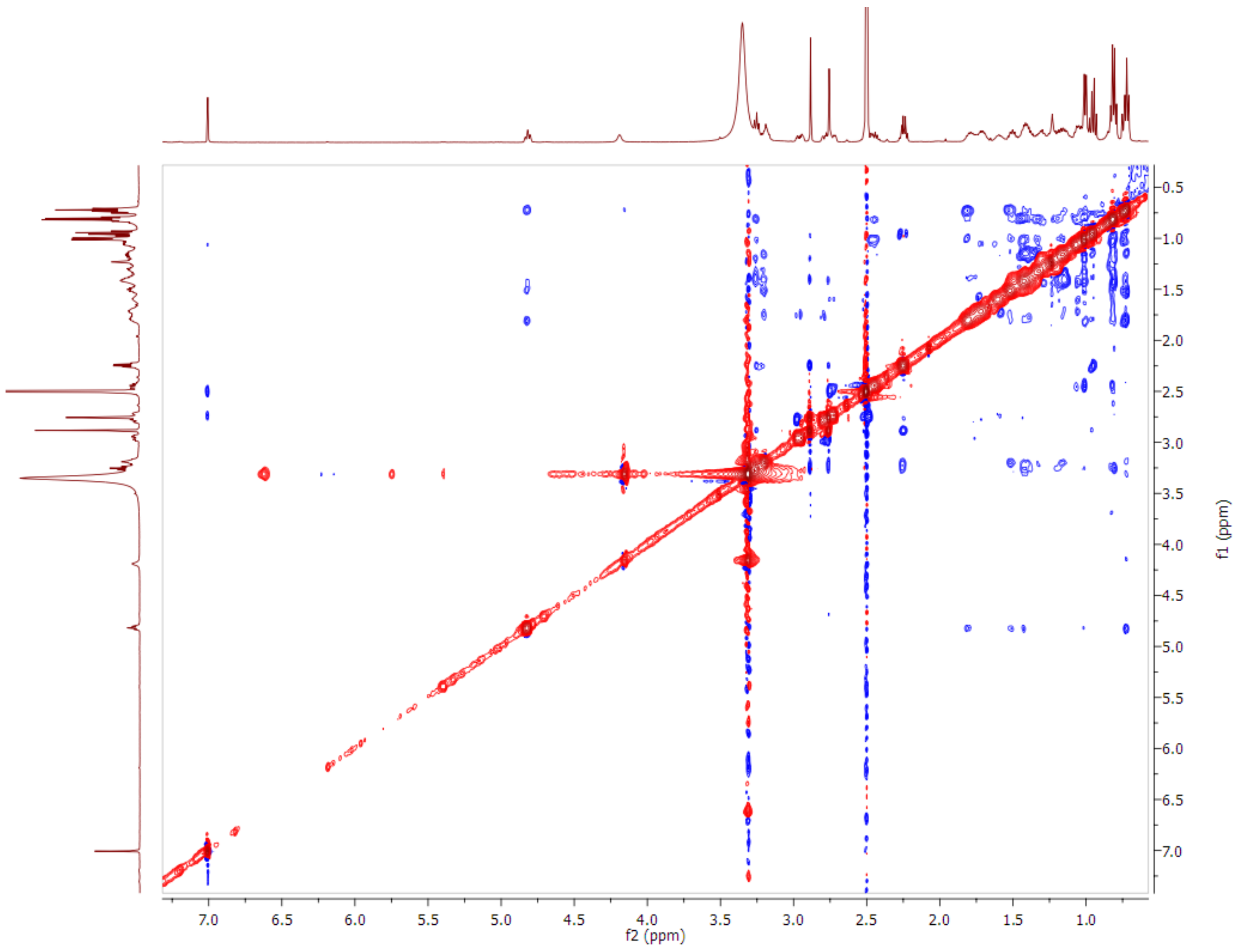


Figure S11. NOESY of **1**.

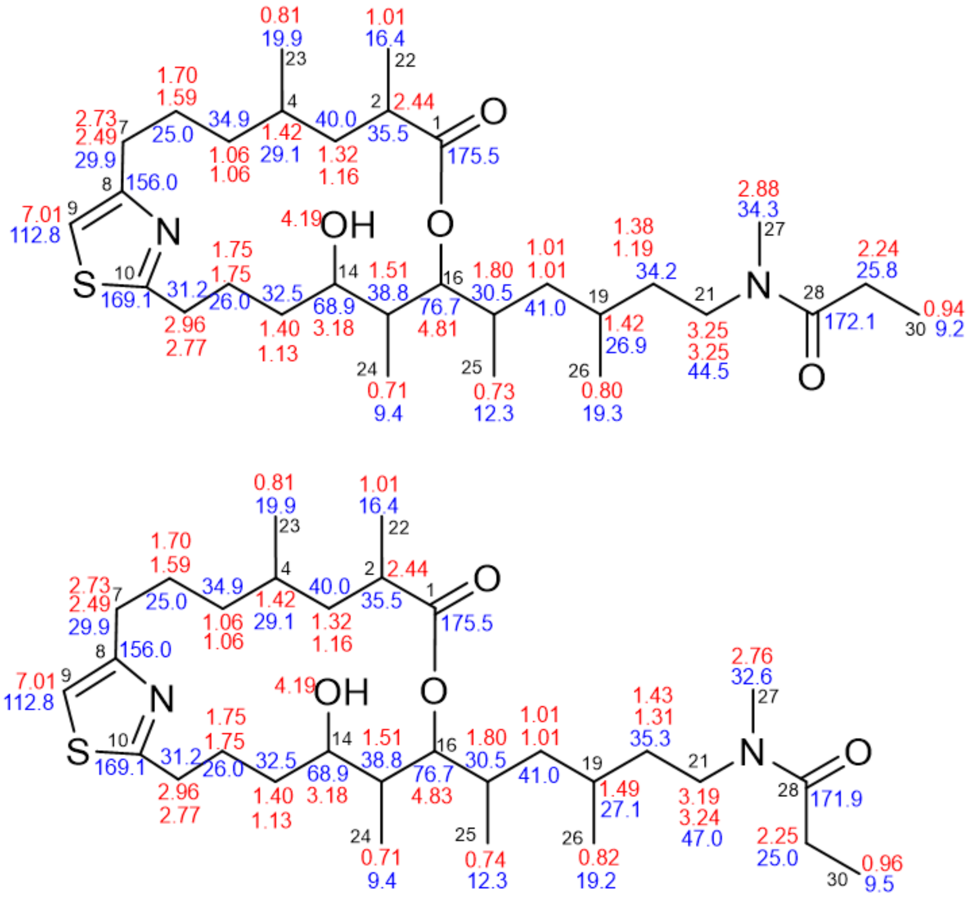


**1**

Figure S12. $^{1}H$ (red) and $^{13}C$ (blue) NMR chemical shifts for the *Z* (top) and *E* (bottom) conformers of **1**.

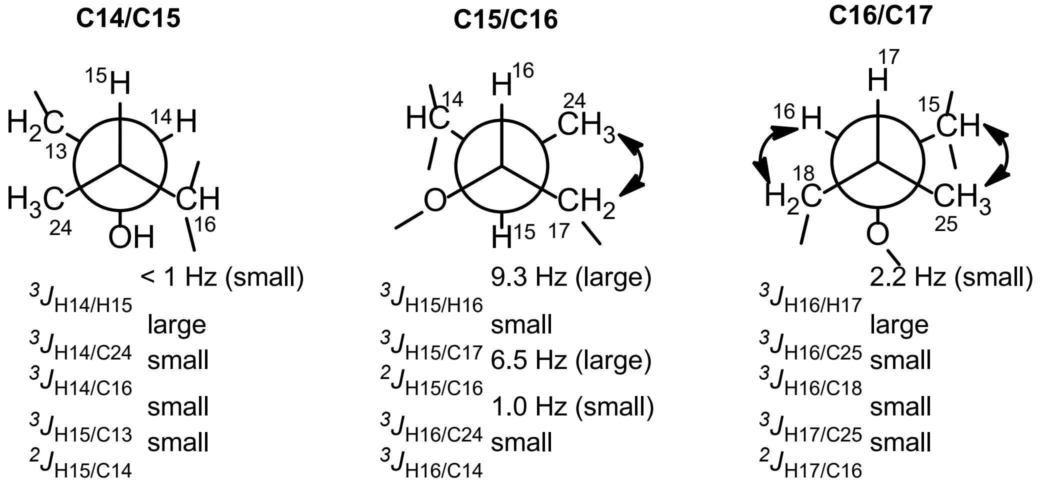


Figure S13. Relative configuration of the of the segment C-14 to C-17 of **1** as determined using the Murata's method.[30] A few $^{2,3}J_{CH}$ were measured using a HECADE-HSQC experiment,[31] but in most cases this was impossible because the small couplings between H-14 and H-15 and between H-16 and H-17 blocked the TOCSY coherence transfer on which the HECADE-HSQC experiment is based. The remaining $^{2,3}J_{CH}$ were estimated as "large" or "small" from the ratio of the relative magnitudes of their HMBC peaks with respect to a common proton. Double-headed arrows represent NOE between the indicated protons.

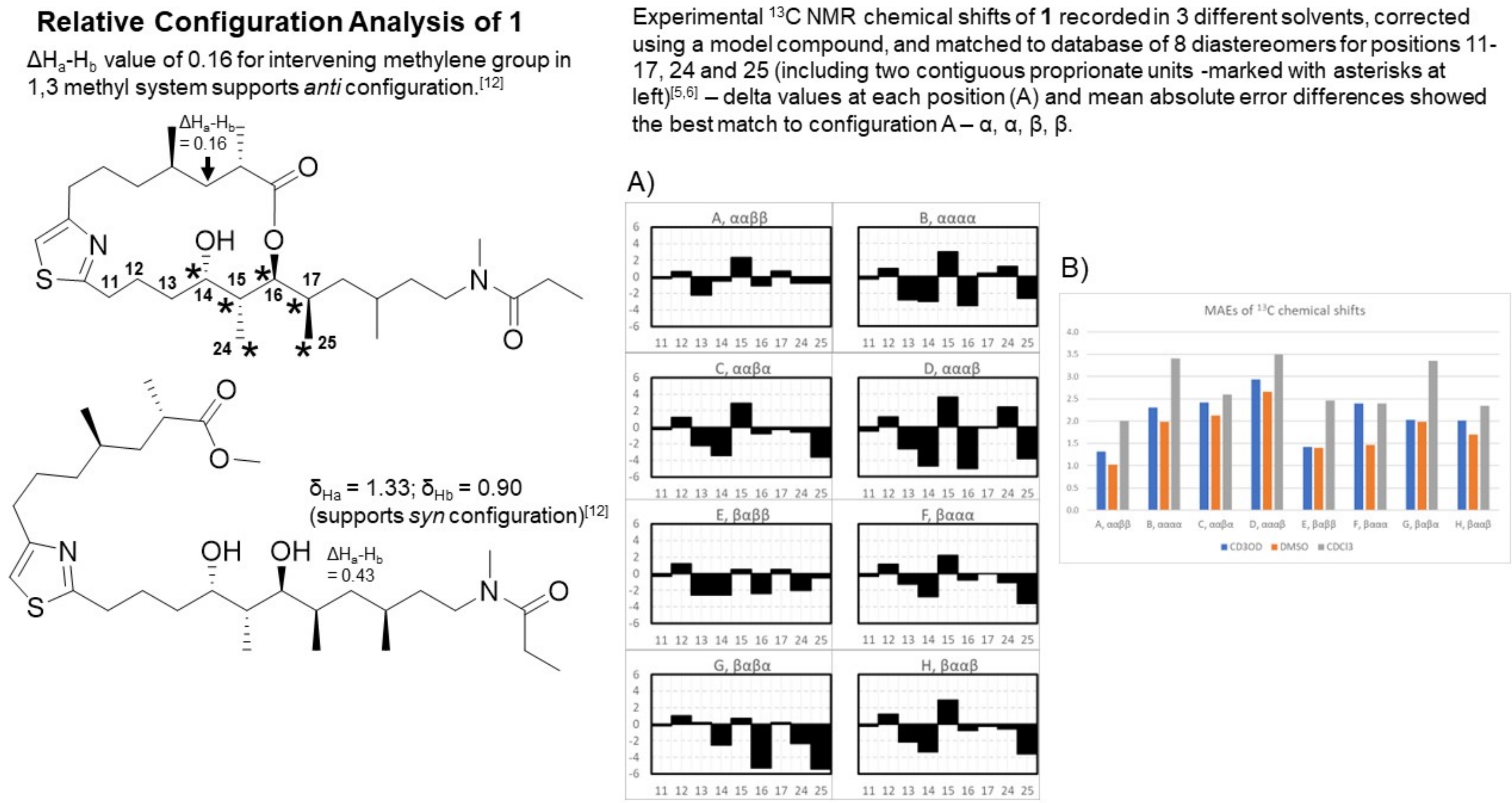


Figure S14. Relative configuration analysis of **1**. Differences in the chemical shifts between diastereotopic protons were used to define the relative configuration of the 1,3-methyl systems in the intact molecule and following methanolysis. The configuration of the contiguous propionate units was determined following NMR analysis and comparison to the NMR database developed by Kobayashi and coworkers.[30]

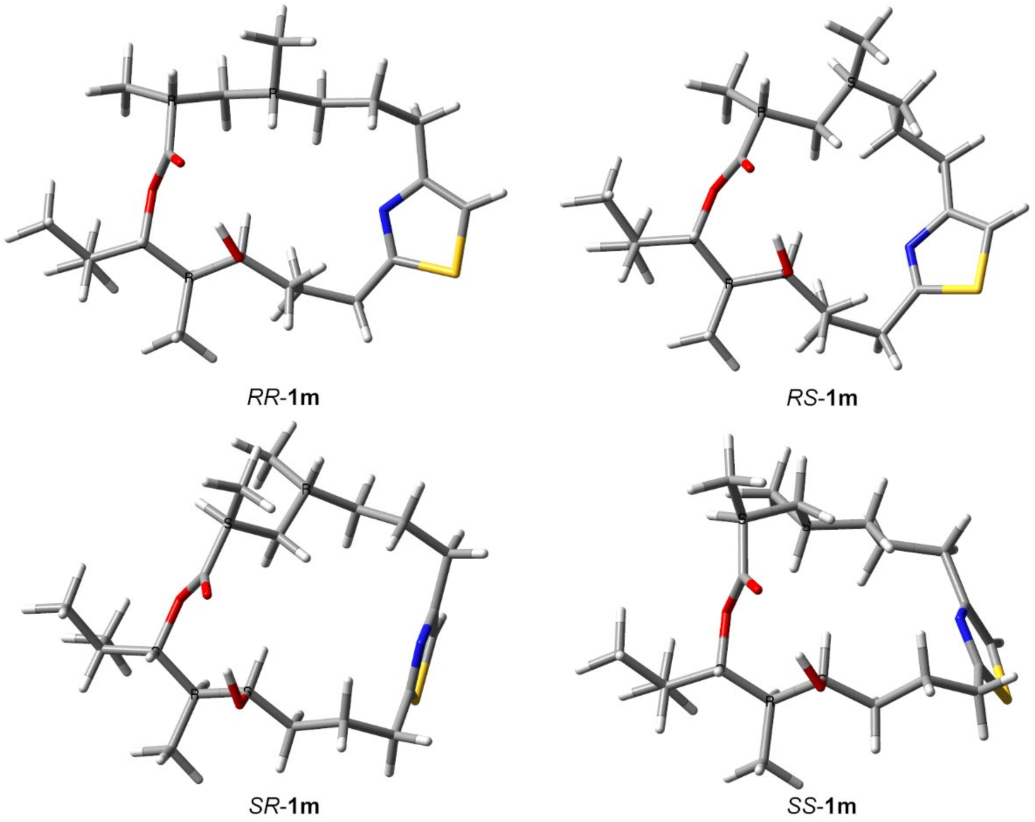


Figure S15. Lowest energy conformation of model compounds *RR*-**1m**, *RS*-**1m**, *SR*-**1m**, *SS*-**1m** at the B3LYP/6-31G(d,p)/SMD(DMSO) level of theory

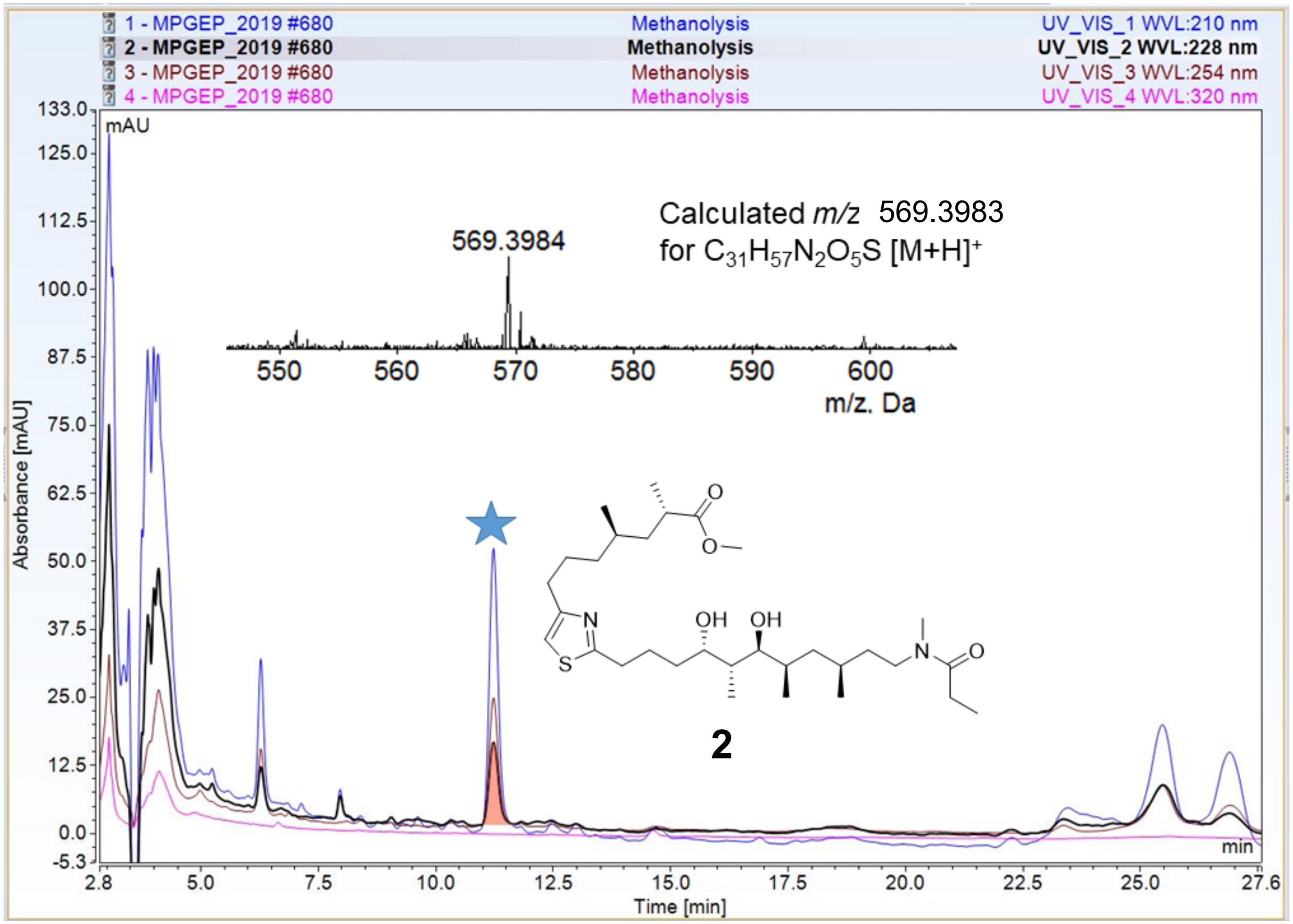


Figure S16. Isolation and HRMS confirmation of the formation of the methyl ester of **1**.

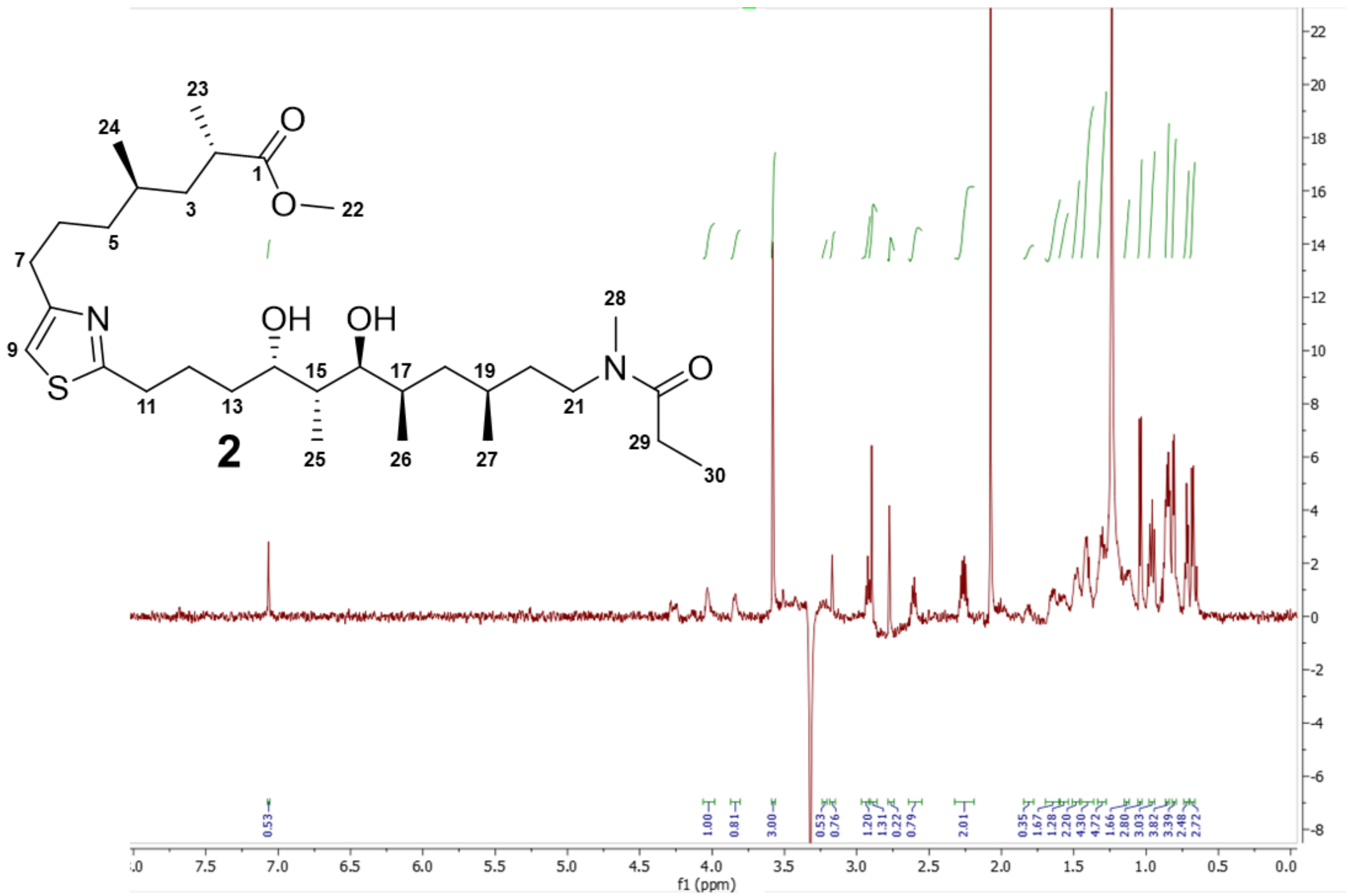


Figure S17. $^{1}H$ NMR of the trichothilone methyl ester (**2**) (600 MHz, DMSO-$d_6$) with solvent suppression.

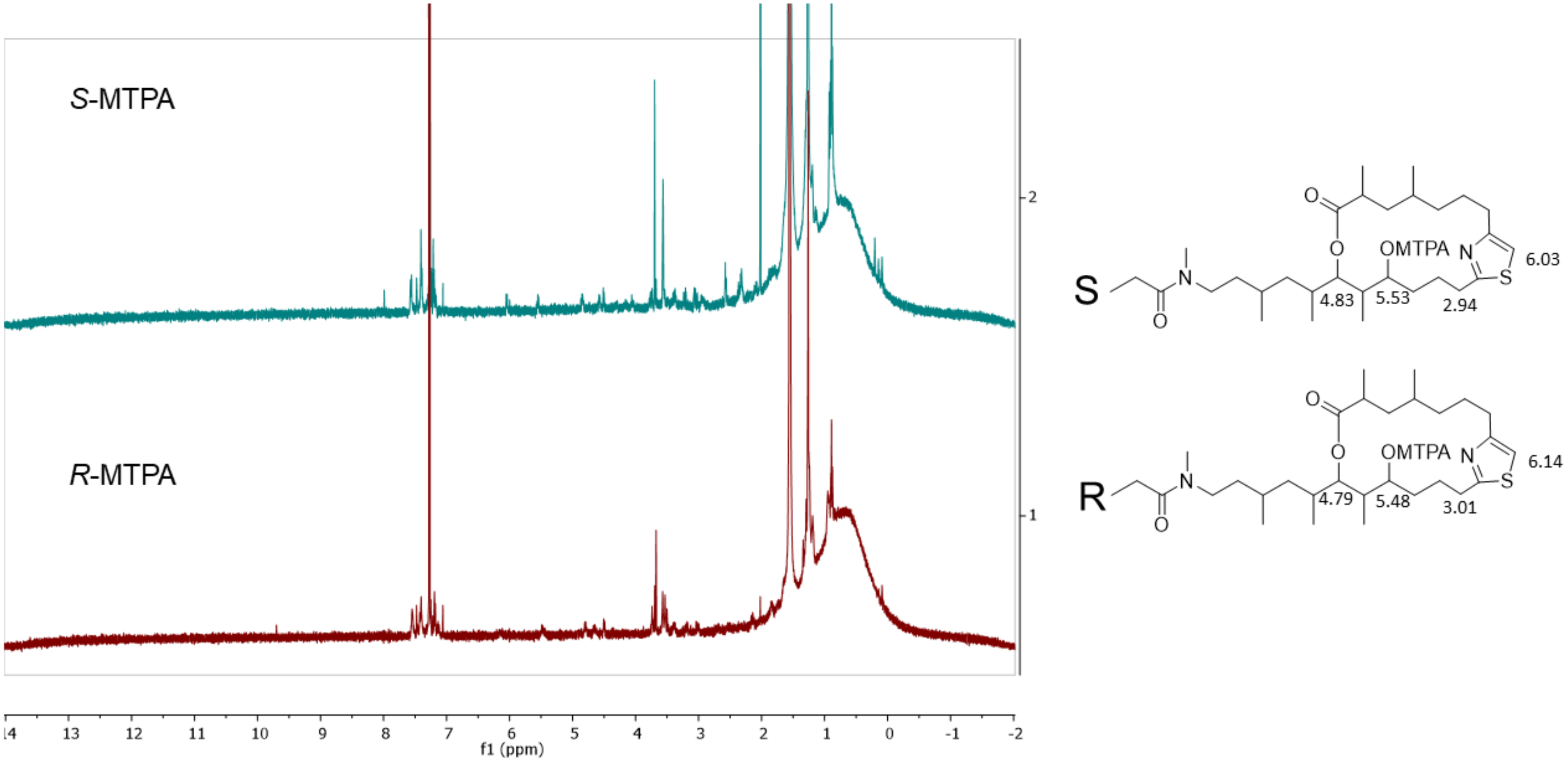


Figure S18. Mosher ester analysis of **1**.

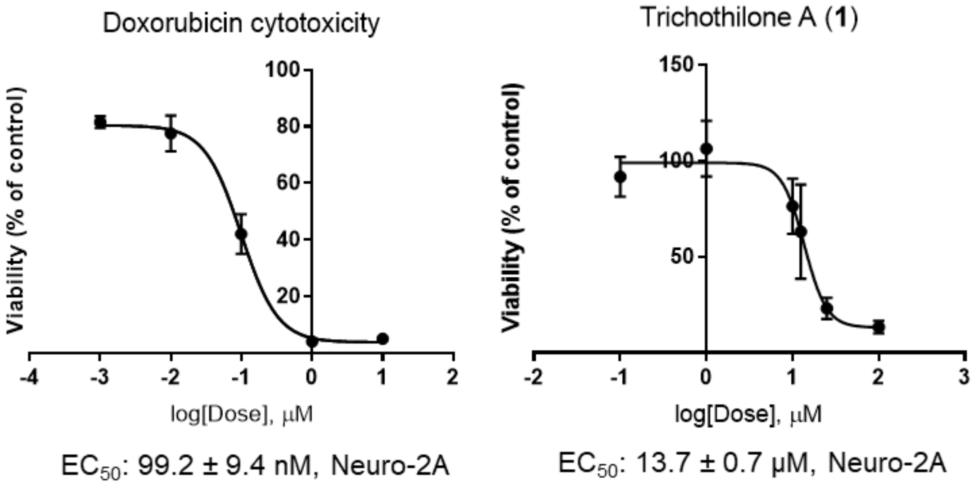


Figure S19. Cytotoxicity of **1** compared to the positive control doxorubicin against neuro-2A murine neuroblastoma cells.

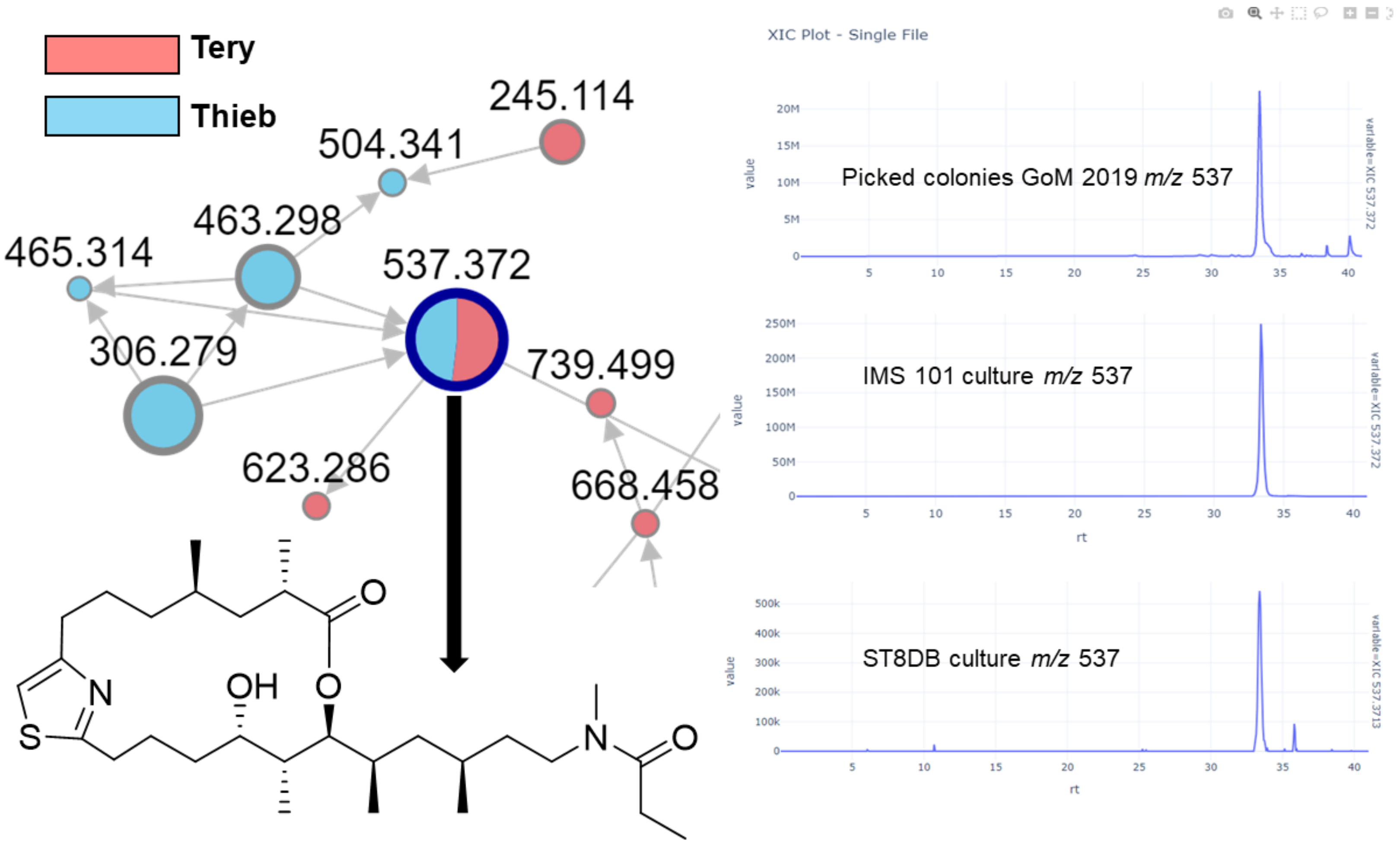


Figure S20. Molecular network of extracts from GoM2019-11 and IMS101 and ST8 showing the trichothilone A (**1**) node present in both species. Extracted ion chromatogram (XIC) shows detection of **1** in *T. erythraeum* IMS101 and ST8 extracts.

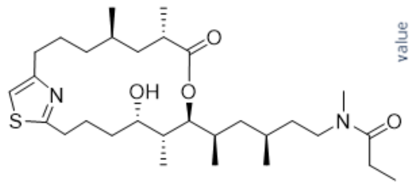


**1**

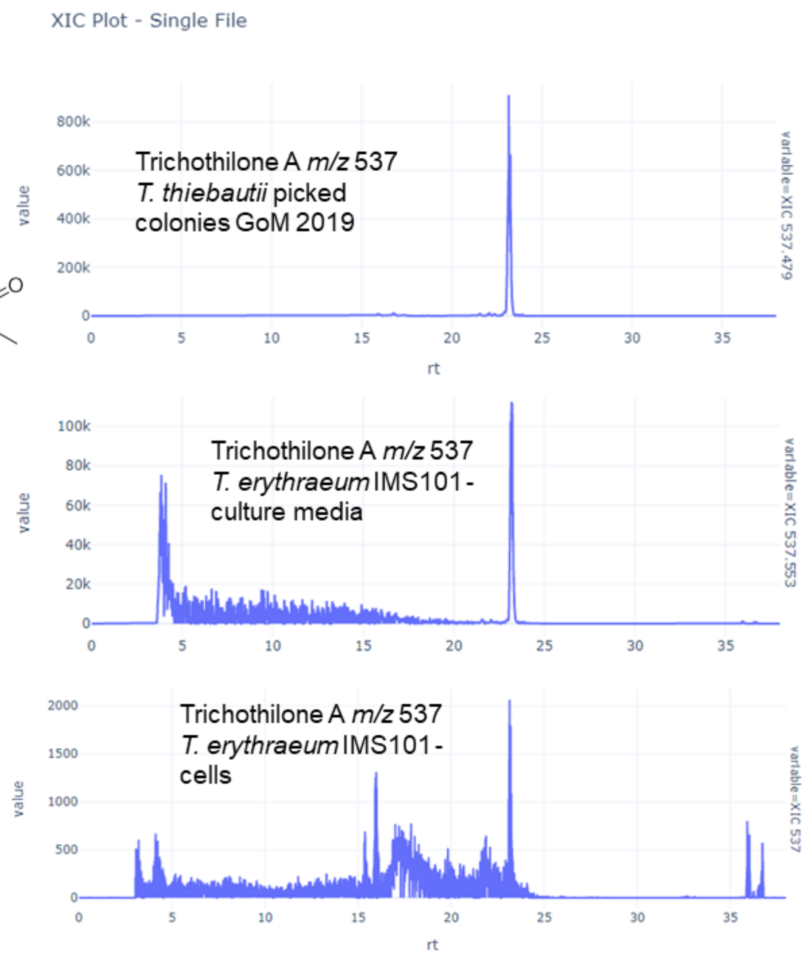


Figure S21. Trichothilone A (**1**) was detected in both the cells and media of *T. erythraeum* culture IMS101.

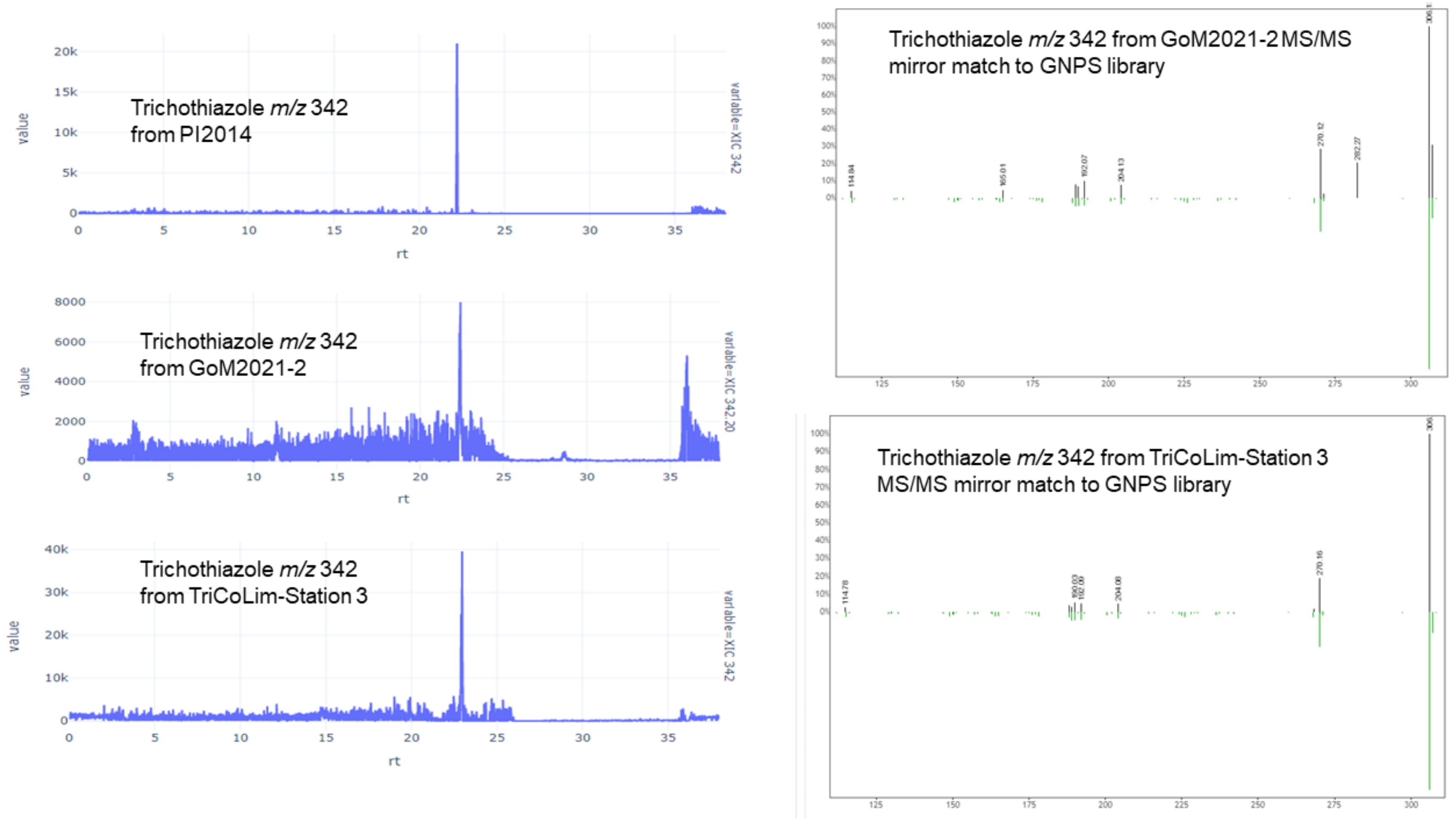


Figure S22. Confirmation of trichothiazole A in samples from GoM2021 and TriCoLim by LC-MS retention time and MS/MS fragmentation.

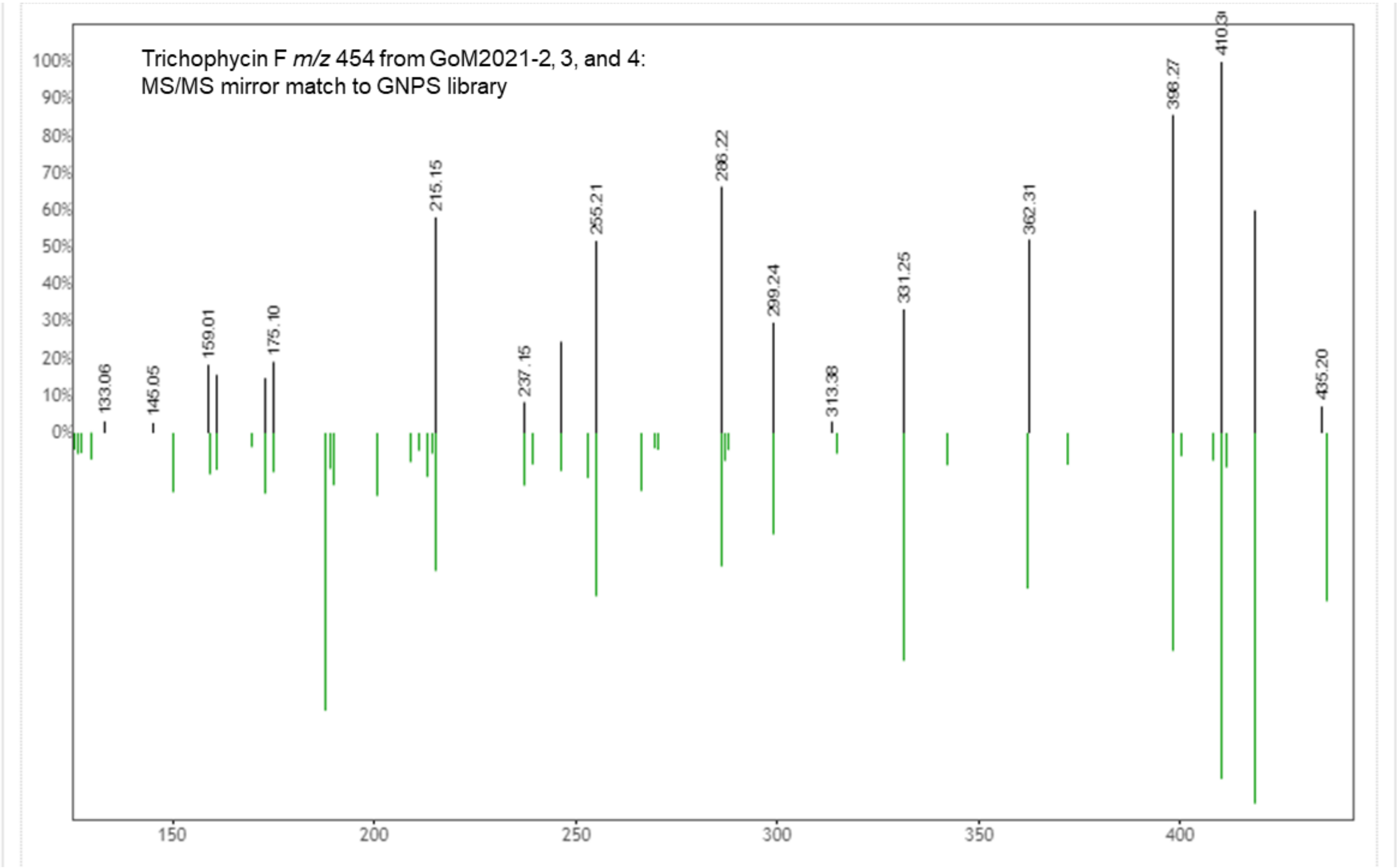


Figure S23. Confirmation of trichophycin F in samples from GoM2021 and TriCoLim by MS/MS fragmentation.

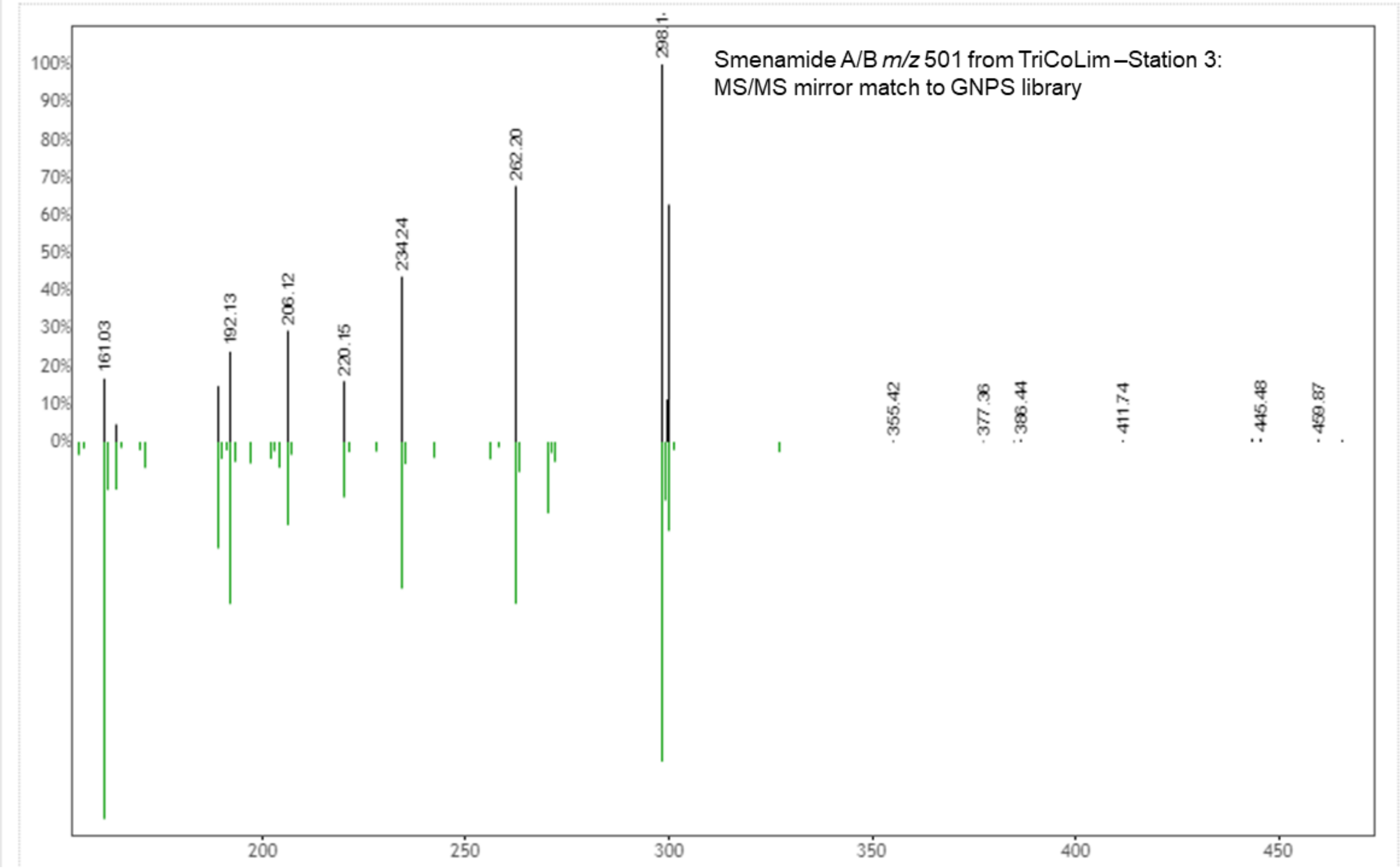


Figure S24. Confirmation of smenamide A/B in samples from GoM2021 and TriCoLim by MS/MS fragmentation.

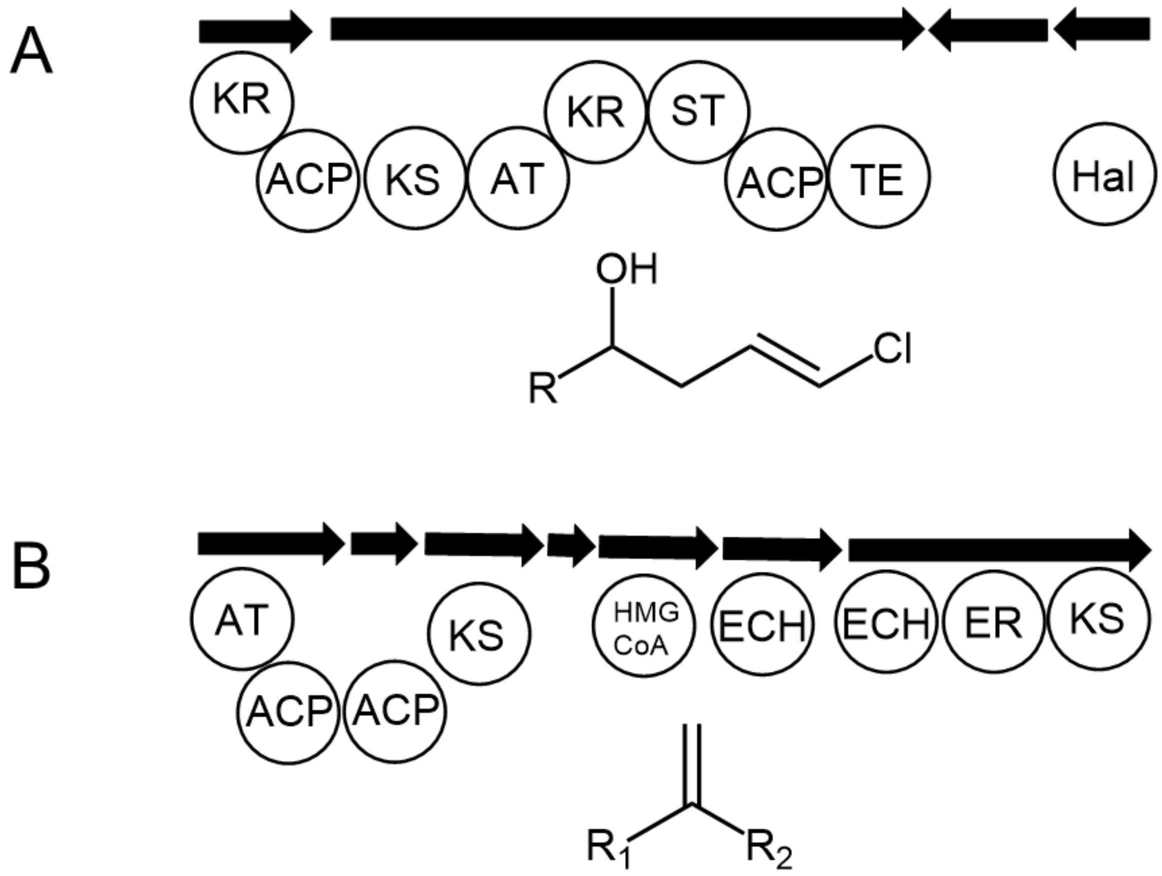


Figure S25. Partial putative biosynthetic gene cluster than encode the terminal vinyl chloride (A) and vinylidene (B) groups in *Trichodesmium* chlorinated metabolites. KS = ketosynthase, AT = acyl transferase, ACP = acyl carrier protein, KR = ketoreductase, ST = sulfotransferase, TE = thioesterase, Hal = halogenase, HMG CoA = hydroxylmethylglutaryl-CoA synthase, ECH = enoyl-CoA hydratase, ER = enoyl reductase

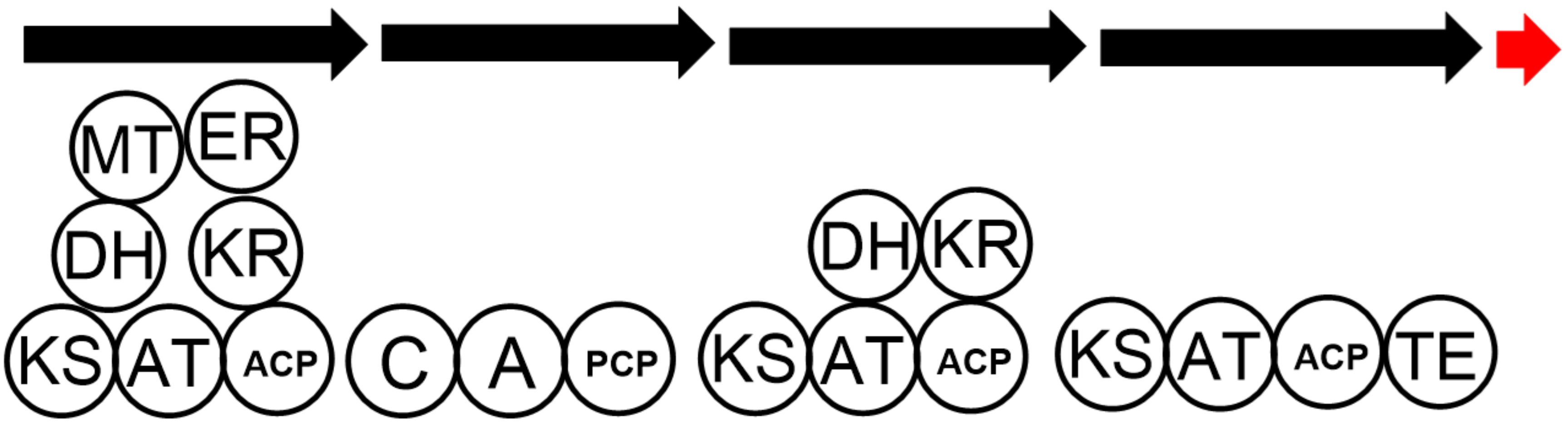


Figure S26. Orphan gene cluster found in multiple *Trichodesmium* MAGs. MT = methyltransferase, ER = enoyl reductase, KS = ketosynthase, AT = acyl transferase, ACP = acyl carrier protein, KR = ketoreductase, DH = dehydratase, C = condensation, A = adenylation, PCP = peptidyl carrier protein, TE = thioesterase. Red arrow illustrates a lipocalin-like gene.

### 4. References


[1] J. Komárek, K. Anagnostidis, Cyanoprokarota Part 2: Oscillatoriales. München, Germany: Elsevier, **2005**.

[2] M. J. Bertin, P. G. Wahome, P. V. Zimba, H. He, P. D. R. Moeller, *Mar. Drugs* **2017a**, *15*, https://doi:10.3390/md15010010.

[3] K. M. McManus, R. D. Kirk, C. W. Via, J. S. Lotti, A. F. Roduit, R. Teta, S. Scarpato, A. Mangoni, M. J. Bertin, *J. Nat. Prod*. **2020**, *83*, 2664-2671. https://doi.org/10.1021/acs.jnatprod.0c00550.

[4] E. A. Webb, N. A. Held, Y. Zhao, E. D. Graham, A. E. Conover, J. Semones, M. D. Lee, Y. Feng, F. Fu, M. A. Saito, D. A. Hutchins, *ISME Commun.* **2023**, *3* (1), 15. https://doi.org/10.1038/s43705-023-00214-y.

[5] Y. Kobayashi, J. Lee, K. Tezuka, Y. Kishi, *Org. Lett.* **1999**, *1* (13), 2177–2180. https://doi.org/10.1021/ol9903786.

[6] J. Lee, Y. Kobayashi, K. Tezuka, Y. Kishi, *Org. Lett.* **1999**, *1* (13), 2181-2184. https://doi.org/10.1021/ol990379y.

[7] K. E. Gilbert, Pcmodel (version 10.0), Serena Software, Bloomington, IN, 2013.

[8] Gaussian 16, Revision C.01, Gaussian Inc., Wallingford CT, USA.

[9] G. K. Pierens, *J. Comput. Chem.* **2014**, *35* (18), 1388–1394. https://doi.org/10.1002/jcc.23638.

[10] N. Grimblat, M. M. Zanardi, A. M. Sarotti, *J. Org. Chem*. **2015**, *80* (24), 12526–12534. https://doi.org/10.1021/acs.joc.5b02396.

[11] M. Wang, J. J. Carver, V. V. Phelan, L. M. Sanchez, N. Garg, Y. Peng, D. D. Nguyen, J. Watrous, C. A. Kapono, T. Luzzatto-Knaan, C. Porto, A. Bouslimani, A. V. Melnik, M. J. Meehan, W.-T. Liu, M. Crüsemann, P. D. Boudreau, E. Esquenazi, M. Sandoval-Calderón, R. D. Kersten, L. A. Pace, R. A. Quinn, K. R. Duncan, C.-C. Hsu, D. J. Floros, R. G. Gavilan, K. Kleigrewe, T. Northen, R. J. Dutton, D. Parrot, E. E. Carlson, B. Aigle, C. F. Michelsen, L. Jelsbak, C. Sohlenkamp, P. Pevzner, A. Edlund, J. McLean, J. Piel, B. T. Murphy, L. Gerwick, C.-C. Liaw, Y.-L. Yang, H.-U. Humpf, M. Maansson, R. A. Keyzers, A. C. Sims, A. R. Johnson, A. M. Sidebottom, B. E. Sedio, A. Klitgaard, C. B. Larson, C. A. Boya P, D. Torres-Mendoza, D. J. Gonzalez, D. B. Silva, L. M. Marques, D. P. Demarque, E. Pociute, E. C. O'Neill, E. Briand, E. J. N. Helfrich, E. A. Granatosky, E. Glukhov, F. Ryffel, H. Houson, H. Mohimani, J. J. Kharbush, Y. Zeng, J. A. Vorholt, K. L. Kurita, P. Charusanti, K. L. McPhail, K. F. Nielsen, L. Vuong, M. Elfeki, M. F. Traxler, N. Engene, N. Koyama, O. B. Vining, R. Baric, R. R. Silva, S. J. Mascuch, S. Tomasi, S.

Jenkins, V. Macherla, T. Hoffman, V. Agarwal, P. G. Williams, J. Dai, R. Neupane, J. Gurr, A. M. C. Rodríguez, A. Lamsa, C. Zhang, K. Dorrestein, B. M. Duggan, J. Almaliti, P.-M. Allard, P. Phapale, L.-F. Nothias, T. Alexandrov, M. Litaudon, J.-L. Wolfender, J. E. Kyle, T. O. Metz, T. Peryea, D.-T. Nguyen, D. VanLeer, P. Shinn, A. Jadhav, R. Müller, K. M. Waters, W. Shi, X. Liu, L. Zhang, R. Knight, P. R. Jensen, B. Ø. Palsson, K. Pogliano, R. G. Linington, M. Gutiérrez, N. P. Lopes, W. H. Gerwick, B. S. Moore, P. C. Dorrestein, N. Bandeira, *Nat. Biotechnol.* **2016**, *34*, 828–837.

[12] Y. Schmidt, K. Lehr, L. Colas, B. Breit, *Chem. Eur. J.* **2012**,18, 7071-7081. https://doi.org/10.1002/chem.201103988.
